# Global threat abatement potential for terrestrial vertebrates

**DOI:** 10.64898/2026.06.12.731583

**Authors:** Francesca A. Ridley, Leon Bennun, Thomas M. Brooks, Stuart H. M. Butchart, Michael W Dales, Frank Hawkins, Randall R. Jiménez, Nicholas B.W. Macfarlane, Philip J. McGowan, Thomas Starnes, Simon Tarr, Joseph A. Turner, Daniele Baisero, Janice Chanson, Neil Cox, Vivek Menon, Kelsey Neam, Michela Pacifici, Andrés Rodríguez, Jon Paul Rodríguez, Carlo Rondinini, Louise Mair

## Abstract

1.

**Aim:** The Species Threat Abatement and Restoration (STAR) metric was developed to support setting and measuring progress towards science-based targets for species conservation, in alignment with the Kunming-Montreal Global Biodiversity Framework. The STAR metric quantifies the potential reduction in species’ global extinction risk achievable through actions to abate threats (STAR_T_) and restore habitat (STAR_R_). The STAR metric is used across multiple sectors to assess contributions to nature-positive species outcomes and implement action for biodiversity. Here we present a substantially enhanced global estimate of STAR_T_ for terrestrial vertebrates. Developed in response to user demand, this work integrates advancements in methodology and data quality, increased spatial resolution, and addition of reptile species.

**Location:** Global

**Time Period:** Current

**Major taxa studied:** Terrestrial vertebrates

**Methods:** STAR_T_ was estimated at 1km^2^ resolution for 9,100 species of threatened and Near Threatened birds, mammals, amphibians and reptiles, using IUCN Red List assessments (version 2025-1) and area of habitat (AOH) maps generated using an advanced data-driven methodology and satellite-derived land-cover data. AOH maps were validated using a two-stage protocol using species observation data.

**Results:** The six countries with the highest estimated STAR_T_ scores, and therefore the largest individual potential to reduce global extinction risk via tackling threats nationally were Brazil, Madagascar, Indonesia, Mexico, Ecuador and Colombia (each contributing over 5% of global estimated STAR_T_). The threat with the greatest individual potential to reduce extinction risk was annual and perennial non-timber crops (21.2% of global STAR_T_).

**Main conclusions:** Targeted actions to tackle a few high-impact threats in a few discrete locations, and cumulative effort across multiple areas with lower individual potential, are both required to meaningfully reduce species’ extinction risk through threat abatement. This global scale estimation of STAR_T_ enables consistent scoping of conservation opportunity over large areas and provides the critical initial data to support planning and action.

## 2. Introduction

In response to ongoing human-driven declines in global biodiversity (IPBES, 2019; CBD, 2020), in 2022 Parties to the Convention on Biological Diversity agreed the Kunming-Montreal Global Biodiversity Framework (KMGBF). The KMGBF includes 4 goals and 23 targets, including the aim to halt human-induced extinction and reduce the extinction risk of known threatened species (CBD/COP/DEC/15/4). The Species Threat Abatement and Restoration (STAR) metric was designed to support the achievement of this goal (Mair et al., 2021) by informing the setting of – and measurement of progress towards – science-based targets for species conservation, and the metric has been adopted as a complementary indicator for Targets 4 (urgent action to halt species extinctions and reduce extinction risk) and 15 (assess, disclose and reduce the impacts of businesses on biodiveristy)(CBD/COP/DEC/15/5).

The STAR metric quantifies the potential benefit to species of actions to abate (reduce and mitigate) threats and restore habitat (Mair et al., 2021). The STAR metric is calculated for all threatened and Near-Threatened species of comprehensively assessed taxonomic groups (i.e. groups for which >80% of species has been assessed on the IUCN Red List of Threatened Species), which at the time of initial publication was terrestrial birds, mammals and amphibians (Mair et al., 2021). In theory, if all threats were successfully abated a species’ extinction risk would be reduced to ‘Least Concern’. Threat abatement potential (STAR_T_) and restoration potential (STAR_R_) represent two distinct, and equally important, components of the STAR metric. Here, we focus on estimating STAR_T_, which represents the potential benefit to species of actions to abate threats in particular locations.

Since its original publication, the STAR metric (Mair et al., 2021) has gained significant traction within the conservation science community, having been cited over 124 times at the time of writing according to Altmetrics. These citations reveal several emerging themes in its application and interpretation. Some authors expand STAR to other biomes (Turner et al., 2024), provide assessments to allow expansion of STAR to include other taxonomic groups, including reptiles (Cox et al., 2022) and plants (Bachman, Brown, Leão, Nic Lughadha, & Walker, 2024), or highlight the need for comparable methods for ecosystems (Nicholson et al., 2024). Additional research has explored the use of STAR for evaluating contributions to the KMGBF on the national (Mair et al., 2023) and regional (Chaudhary et al., 2022; Guerrero-Pineda et al., 2022; Jiménez et al., 2025) scale, and threat abatement potential attributed to international trade (Irwin & Geschke, 2023; Irwin et al., 2022; Irwin, Geschke, & Mackenbach, 2024).

Another common theme is the use of STAR for business-related disclosures and to measure progress toward nature positive outcomes (Hawkins et al., 2024; Layman et al., 2024; Zhu, Prescott, Chu, & Carrasco, 2024; Zu Ermgassen et al., 2022). STAR is the central metric in IUCN’s Rapid High-Integrity Nature-positive Outcomes (RHINO) approach (IUCN, 2025) which was designed to align with the LEAP (Locate, Evaluate, Assess, and Prepare) approach of the Taskforce on Nature-related Financial Disclosures (TNFD, 2024). The STAR metric is served for commercial use through the Integrated Biodiversity Assessment Tool (IBAT; https://www.ibat-alliance.org). As such, our further development of the STAR metric is timely to meet real-world user needs and demand.

At the time of writing there have been 14 IUCN Red List updates since the data used in Mair et al. (2021), reflecting revisions in taxonomy and new assessments of species extinction risk, with some species being reassessed and others being assessed for the first time. The use of an updated version of the Red List (IUCN, 2025) means that data for an additional 3,741 threatened and Near Threatened species can be included. These represent some new species of previously included taxonomic groups (either newly assessed or previously Least Concern or Data Deficient) and an expansion in taxonomic scope to incorporate reptiles which are now comprehensively assessed (Cox et al., 2022).

There have also been several advancements in the methods for calculating area of habitat (AOH). AOH is the area within a species’ geographic range containing suitable land-cover at suitable elevation (Brooks et al., 2019). For STAR_T_, a species’ current AOH is used as the basis for estimating the contribution of actions in a particular area of interest to the species’ global extinction risk, assuming that populations are evenly distributed throughout each species’ AOH. In the original global STAR estimation (Mair et al., 2021), AOH was mapped at 5 x 5 km resolution. There is an increasing user-driven demand for higher-resolution data to improve spatial precision of STAR assessments and to better inform local and landscape-level conservation planning.

Calculating a species’ AOH relies on translating the thematic habitat classes from the IUCN Habitats Classification Scheme (version 3.1) (IUCN, 2012a) to spatially explicit land-cover classes (also termed a ‘crosswalk’). In Mair et al. (2021) this crosswalk was based on the thematic similarity of habitat class descriptions to land-cover descriptions and was established by ecological experts. Since then, a data-driven approach has been developed and tested on terrestrial birds and mammals, in which the association between habitat classes of the IUCN Habitat Classification Scheme and land-cover is modelled using species occurrence data (Lumbierres, Dahal, Di Marco, et al., 2022). The use of a data-driven approach has several benefits including removal of researcher bias and allowing calculation of uncertainty.

New and more accurate digital surface models (DSMs) are also available for AOH mapping. The new Copernicus GLO-30 DSM, which is based on TanDEM-X data, is considered the most recent and accurate elevation dataset (Guth & Geoffroy, 2021). Hawker et al. (2022) further applied a machine learning correction algorithm to remove forests and buildings to generate the best available global representation of ground elevation. Updating the global estimate of species threat abatement potential using these improved inputs is likely to enhance the accuracy of AOH maps significantly and refine our understanding of where actions to abate threats could have the greatest benefit for species.

Our aim was to improve global understanding of the reduction in terrestrial vertebrate extinction risk that could be achieved through actions to abate threats. We achieved this by first evaluating whether the advancement in AOH mapping methodology, that was developed using data on birds and mammals (Dahal, Lumbierres, Butchart, Donald, & Rondinini, 2022; Lumbierres, Dahal, Di Marco, et al., 2022; Lumbierres, Dahal, Soria, et al., 2022), can be applied to map suitable habitat for amphibians and reptiles. We then mapped AOH at increased spatial resolution (now 1 km^2^) and calculated STAR_T_ using updated assessments of species extinction risk from the IUCN Red List of Threatened Species (version 2025-1). We specifically sought answers to the following questions: (1) what is the global distribution of threat abatement potential for terrestrial vertebrates, (2) which threats to terrestrial vertebrates have the greatest abatement potential for each major geographic region and (3) how do these distributions of threat abatement potential differ from those found in Mair et al. (2021).

## 3. Methods

### 3.1 Species data

Species’ taxonomy, habitat associations, identified threats, elevation limits, and range data were downloaded from the IUCN Red List of Threatened Species (version 2025-1, hereafter simply ‘Red List’) (IUCN, 2025) for all terrestrial Near Threatened or threatened (i.e. Vulnerable, Endangered or Critically Endangered) bird, mammal, amphibian and reptile species. Subspecies and sub-populations were not included. The downloaded data included assessments of extinction risk for 9,586 species, including 2,217 birds, 1,718 mammals, 3,264 amphibians and 2,387 reptiles, on which further filtering was applied on species’ threat and range polygon attributes.

Species’ threats were refined to those with some current or expected future impact. Species’ threats were removed if the timing was ‘past, and unlikely to return’. Threat scope and severity scores were combined into an estimated percentage population decline according to Table S1. Where either or both of scope and severity were missing or unknown for any threat (27,826 species-threat combinations, 66%; See Appendix S1 in the supporting information), the scope was assumed to be ‘Majority’ and severity was assumed to be ‘Slow, significant Declines’ following Mair et al. (2021). As scope and severity are only required where assessors have listed both major and minor threats (IUCN, 2012b), we assume that where scope and severity are absent the impact to the species is the same. Species’ threats with zero expected population decline were removed, resulting in the removal of species where the only reported threats had a negligible impact. Threats were processed at the level of specificity reported in the Red List Assessments with each named invasive alien species or virus considered a distinct threat, following Mair et al. (2021). A total of 9,350 species had at least one relevant threat, including 2,143 birds, 1,685 mammals, 3,214 amphibians and 2,308 reptiles.

Species’ range polygons, coded according to presence, origin, and seasonality, were extracted from BirdLife International and Handbook of the Birds of the World (2025) for birds, and from the Red List (version 2025-1) for mammals, amphibians and reptiles. All range polygons were included if the species’ presence was recorded as extant or probably extant, and the origin was native, reintroduced or assisted colonisation (i.e. excluding introduced, vagrant and uncertain). Additionally, polygons where species’ presence was possibly extinct were included if the species itself was globally Possibly Extinct or Possibly Extinct in the Wild, as such polygons represent the only plausible range limits for these species (n=278). Species were excluded if their range polygons were generalised by assessors to avoid releasing precise location information for sensitive species (following Red List mapping standards; IUCN, 2021). A total of 222 species had at least one relevant threat but did not have any relevant range polygons and were excluded, including 35 species that had generalised range polygons only or where data on whether the range was generalised was missing. This left 9,128 species for which the AOH mapping methodology was applied.

### 3.2 Calculating species’ Area of Habitat (AOH)

Species’ AOH was calculated by cross-referencing species’ habitat and elevation associations reported in Red List assessments with a crosswalk and spatial datasets on land-cover and elevation. We used the crosswalk developed by Lumbierres, Dahal, Di Marco, et al. (2022), that relates terrestrial habitats at level 1 in the IUCN habitats classification scheme (version 3.1) to at least one discrete land-cover class in the Copernicus Global Land Service Land Cover dataset (CGLS-LC100, version 3.01, 2019 epoch). We used the highest threshold of association between IUCN habitats and CGLS-LC100, as described in Lumbierres, Dahal, Soria, et al. (2022) (Table 1).

**Table 1.**
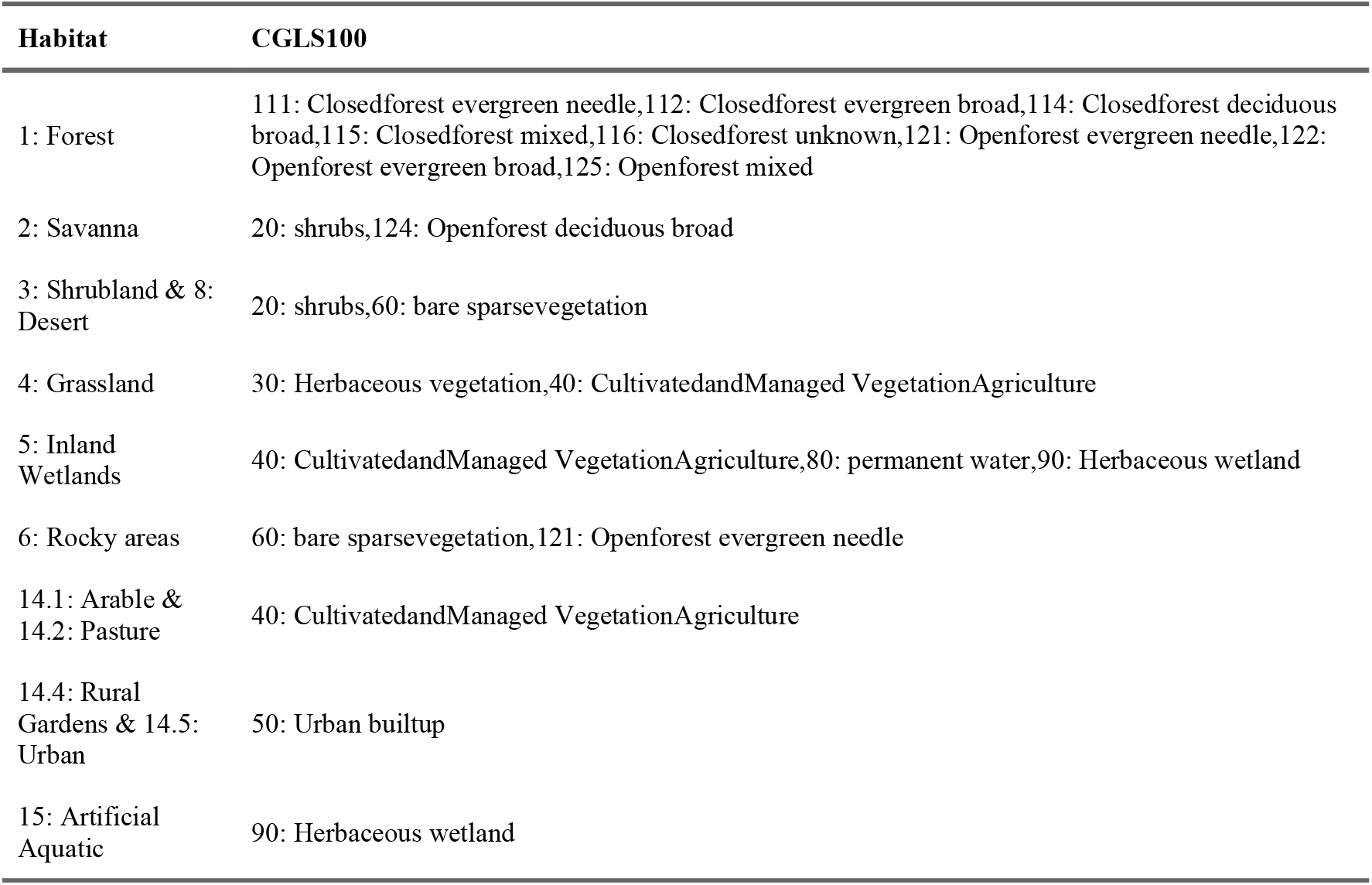
The crosswalk table translating thematic habitat classes at level 1 in the IUCN habitat classification scheme to spatially explicit classes in the Copernicus Global Land Service Land Cover dataset (CGLS100). This Table is reformatted from Lumbierres, Dahal, Di Marco, et al. (2022) and an error was corrected, in which class ‘112: Closedforest evergreen broad leaf’ was incorrectly labelled ‘113: Closed forest evergreen needle leaf’.

| Habitat | CGLS100 |
| --- | --- |
| 1: Forest | 111: Closedforest evergreen needle,112: Closedforest evergreen broad,114: Closedforest deciduous broad,115: Closedforest mixed,116: Closedforest unknown,121: Openforest evergreen needle,122: Openforest evergreen broad,125: Openforest mixed |
| 2: Savanna | 20: shrubs,124: Openforest deciduous broad |
| 3: Shrubland & 8: Desert | 20: shrubs,60: bare sparsevegetation |
| 4: Grassland | 30: Herbaceous vegetation,40: CultivatedandManaged VegetationAgriculture |
| 5: Inland Wetlands | 40: CultivatedandManaged VegetationAgriculture,80: permanent water,90: Herbaceous wetland |
| 6: Rocky areas | 60: bare sparsevegetation,121: Openforest evergreen needle |
| 14.1: Arable & 14.2: Pasture | 40: CultivatedandManaged VegetationAgriculture |
| 14.4: Rural Gardens & 14.5: Urban | 50: Urban builtup |
| 15: Artificial Aquatic | 90: Herbaceous wetland |

The CGLS-LC100 data were downloaded as a single layer of discrete classes at 100 m resolution. The downloaded layer was reclassified (creating a binary layer per class), reprojected to the Mollweide equal area projection system using the nearest neighbour method then aggregated using the mean, to get the proportional cover of each land-cover class in each 992.2927 x 992.2927 m (hereafter referred to as 1 km^2^) cell. The proportional cover of all relevant land-cover classes was summed per cell to derive the total proportion of each cell that was covered by suitable land-cover classes for a particular species.

Suitable habitat was restricted to areas within a species’ terrestrial range to represent the terrestrial biome only (defined by the presence of any CGLS-LC100 terrestrial land-cover class). To achieve this, two complementary raster layers were produced using the Copernicus discrete land classification data. The class ‘200: Open sea’ was used to represent marine areas and all other classes were considered to represent terrestrial areas, with the exception of cells classed as ‘0: no input data’ that were excluded. Cells at 1 km^2^ resolution that were at least 50% covered by terrestrial land-cover were considered terrestrial. If no suitable land-cover at suitable elevation was identified within the terrestrial range (i.e. “true AOH”), adjustments to the AOH were applied (see Appendix S1). An error in the CGLS100 dataset meant that no terrestrial land-cover classes occurred for a number of island states, overseas territories and offshore islands or islets. A correction to AOH was applied to affected species, in which the terrestrial range and AOH was modified to include affected terrestrial administrative zones (see Appendix S1), but still no terrestrial habitat within the range was able to be located for 28 species, which were therefore excluded from STAR_T_.

The Forests and Buildings Removed Digital Elevation Model (FABDEM) (Hawker et al., 2022), based on the Copernicus GLO 30 DSM (CGLO), and species’ elevation associations in Red List assessments were used to refine the AOH maps to areas of suitable elevation. The generation of FABDEM preceded the most recent version of CGLO30, which contains previously restricted data over Armenia, Azerbaijan and Moldova. We used the updated CGLO data to fill the missing elevation values in this area (40.7371 to 50.9321 degrees longitude and 37.9296 to 45.7696 degrees latitude). The FABDEM was reprojected and aggregated to produce two layers; one representing the minimum and one representing the maximum elevation per 1 km^2^ in the Mollweide equal area projection system. These pre-processed layers are accessible (https://zenodo.org/records/17913105) to promote standardised calculation of AOH. Cells were included in AOH if the elevation range of the cell overlapped the elevation range of the species. Errors and omissions in species’ elevation associations were modified following Baisero (2021) and, for birds, supplemented with ‘occasional’ limits from BirdLife International (2024) (see Appendix S1).

### 3.3 AOH map validation

AOH maps were validated following the two-stage method of Dahal et al. (2022). In step one, a logistic regression model predicting map prevalence was fitted to identify systematic errors in AOH map creation. Map prevalence is the proportion of the range estimated to contain AOH. The fitted predictors consisted of elevation range (meters), elevation mid-point (meters), and the number of habitats (at level 2 in the IUCN Habitats Classification Scheme). All predictors were standardised for inclusion in the model. Different models were fitted for each taxonomic class (birds, mammals, amphibians, reptiles) and a random effect of family was used to account for other unexplained taxonomic variation. The difference between the observed and expected map prevalence values was used to identify and investigate outliers. The Tukey’s fence method of outlier detection was used, where values were considered outliers if they were more than 1.5 times the interquartile range above the third or below the first quartiles (Tukey, 1977).

In step two, AOH maps were validated based on the difference between map prevalence and point prevalence. Point prevalence is the proportion of species’ occurrence observations where fractional coverage of AOH was greater than zero. Species’ point occurrence data from the IUCN Red List (version 2023-1), were supplemented with occurrences from the Global Biodiversity Information Facility (GBIF, see Appendix S1). The distance-weighted, average fractional coverage of AOH surrounding each species point locality was determined using bilinear interpolation of the four nearest cells. The distance-weighting meant that, for a point positioned at the centre of a cell, the fractional cover of AOH around the point was equal to the fractional cover of AOH in the cell. This was used instead of the 300 m buffer applied by Dahal et al. (2022) to better account for positional uncertainty and where points were positioned near the boundary between cells. Species’ AOH maps where point prevalence was greater than map prevalence indicated maps that performed better than expected if AOH were distributed within the species’ range randomly.

The results of both steps were used to identify potential systematic errors in the input data or application of the method (see Appendix S2). Errors were either corrected iteratively or reported to the relevant Red List unit. No species were excluded based on the results of map validation.

Validation results for each species’ AOH map generated, including the number of points available, are available in the Supplementary data.

### 3.4 Calculating threat abatement potential (STAR_T_)

Species threat abatement potential (STAR_T_) was calculated for 9,100 species according to Mair et al. (2021) for country *i* and threat *t* as:

(Equation 1, Mair et al., 2021)

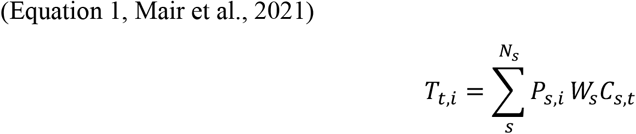

where *P*_s,i_ is the AOH of each species *s* within location *i* (expressed as a percentage of the global species’ current AOH), *W*_s_ is weighting of species *s* based on its Red List extinction risk category (Near Threatened = 1, Vulnerable = 2, Endangered = 3, Critically Endangered = 4), *C*_s,t_ is the relative contribution of threat *t* to the extinction risk of species *s*, and *N*_s_ is the total number of species at location *i* (e.g. each country or 1 km^2^ cell). *C*_s,t_ is the expected population decline (determined by the scope and severity scores) of threat *t*, on species *s*, as a proportion of the total expected population decline of all threats to species *s*. Therefore, the contribution of any one threat to any one species is relative to the contributions of other threats affecting the same species. The STAR_T_ values for a particular location were summed across threats for all Near Threatened and threatened species.

STAR_T_ was calculated for each 1 km^2^ cell in the global distribution in Mollweide equal area projection system. To calculate the difference in threat abatement potential between STAR_T_v1 (Mair et al., 2021) and STAR_T_v2 (described here), we first aggregated STAR_T_ scores from STAR_T_v2 to the 5 x 5 km resolution of STAR_T_v1 by taking the sum. We then calculated the change per 5 x 5 km cell where positive values indicated a higher score in STAR_T_v2 than STAR_T_v1. Methodological differences between STAR_T_v1 and STAR_T_v2 are summarised in Table 2.

**Table 2.**
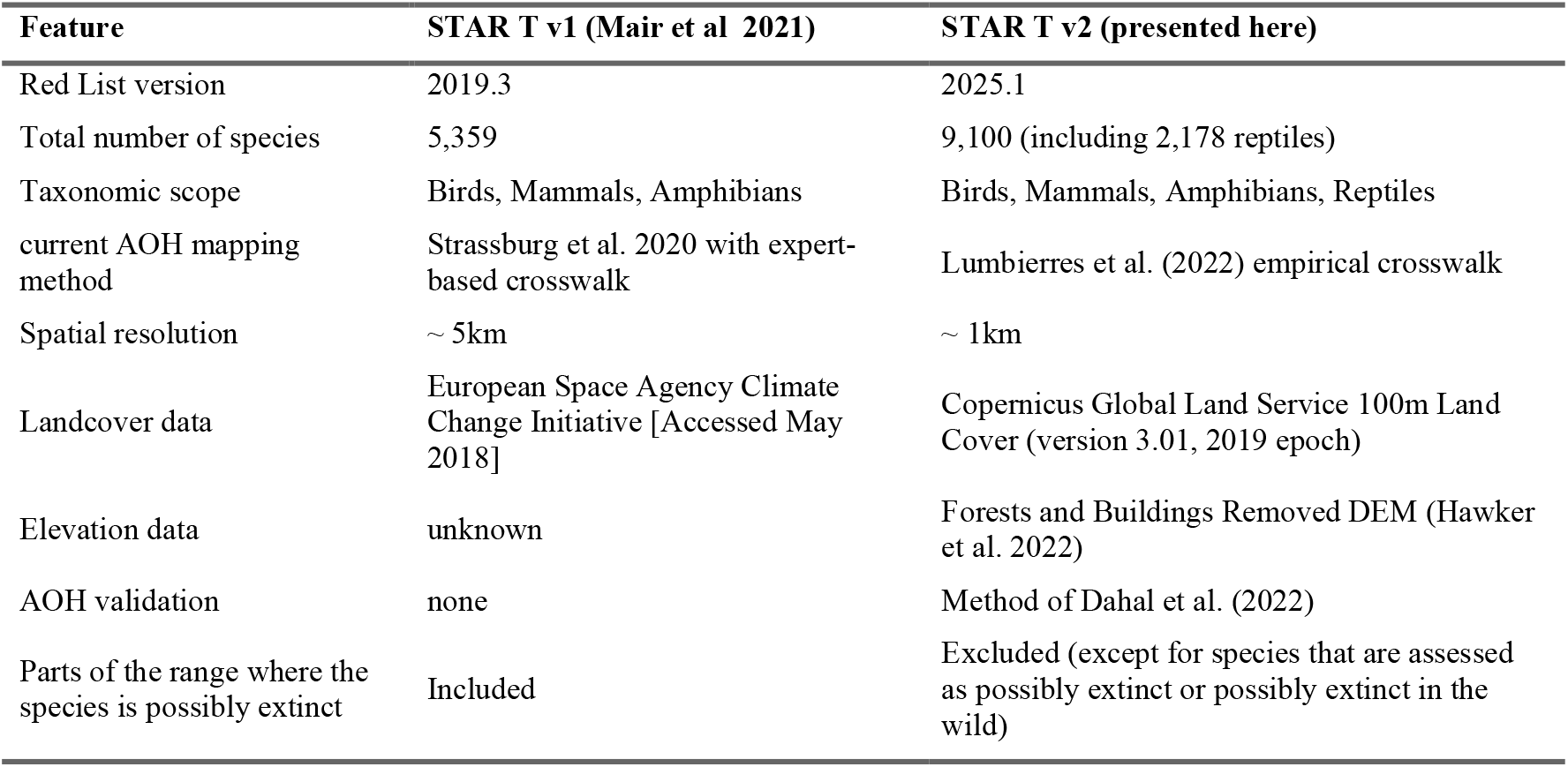
Summary of the methodological differences between STAR_T_v1 (Mair et al., 2021) and STAR_T_v2 (presented here).

| Feature | STAR T v1 (Mair et al 2021) | STAR T v2 (presented here) |
| --- | --- | --- |
| Red List version | 2019.3 | 2025.1 |
| Total number of species | 5,359 | 9,100 (including 2,178 reptiles) |
| Taxonomic scope | Birds, Mammals, Amphibians | Birds, Mammals, Amphibians, Reptiles |
| current AOH mapping method | Strassburg et al. 2020 with expert-based crosswalk | Lumbierres et al. (2022) empirical crosswalk |
| Spatial resolution | ~ 5km | ~ 1km |
| Landcover data | European Space Agency Climate Change Initiative [Accessed May 2018] | Copernicus Global Land Service 100m Land Cover (version 3.01, 2019 epoch) |
| Elevation data | unknown | Forests and Buildings Removed DEM (Hawker et al. 2022) |
| AOH validation | none | Method of Dahal et al. (2022) |
| Parts of the range where the species is possibly extinct | Included | Excluded (except for species that are assessed as possibly extinct or possibly extinct in the wild) |

To calculate STAR_T_ scores per country, polygons from the Global Administrative Areas dataset (GADM, version 4.1)(GADM, 2018) were used to delineate administrative boundaries. Here, the contribution of each country to each species’ extinction risk (*P*_*s,i*_) was the proportion of each species’ global AOH that intersected each country polygon. To reduce unnecessary computation, STAR_T_ scores were only generated for each country/territory where the species was reported to be present in its Red List assessment (applying the same filtering as the IUCN range polygons). Macao and Hong Kong are distinct territories in the Red List data (which follows the ISO-3166 standard) but are sub-regions of China in the GADM data. As such species with AOH in Macao or Hong Kong also contributed to the score of China. Countries were grouped into major geographic regions according to M49 standard codes (UNSD, 1999).

All data processing and analyses were performed in R programming Language (version 4.5.0). R code and final outputs for non-commercial use are accessible in supplementary data. Final outputs for commercial use must be accessed via the Integrated Biodiversity Assessment Tool (IBAT; https://www.ibat-alliance.org).

## 4. Results

### 4.1 Species’ area of habitat mapping and validation

AOH maps were produced for 9,128 species, comprising 2,094 birds, 1,667 mammals, 3,169 amphibians and 2,198 reptiles. These totals include 564 species for which AOH was adjusted or corrections were applied (See Appendix S1 and Appendix S2) and 28 species that were excluded from STAR_T_ because no terrestrial range could be located, even after corrections were applied (Fig. 1). Across all species, mammals had the largest median range size (45,567 km^2^) followed by birds (37,924 km^2^), reptiles (1,780 km^2^) and then amphibians (1,058 km^2^). Among the 8,693 species with a true AOH map (i.e. suitable habitat at suitable elevation within the range), the mean map prevalence (proportion of species’ range that was AOH) was 0.61 (SD 0.27) overall and 0.6 (SD 0.27) for birds, 0.65 (SD 0.26) for mammals, 0.64 (SD 0.26) for amphibians and 0.52 (SD 0.29) for reptiles. For 141 species, the AOH was < 1% of the species’ terrestrial range, yet the difference between observed and predicted map prevalence values identified only 16 species to be outliers (1 birds, 2 mammals, 13 amphibians and 0 reptiles) (see Appendix S2).

**Figure 1.**
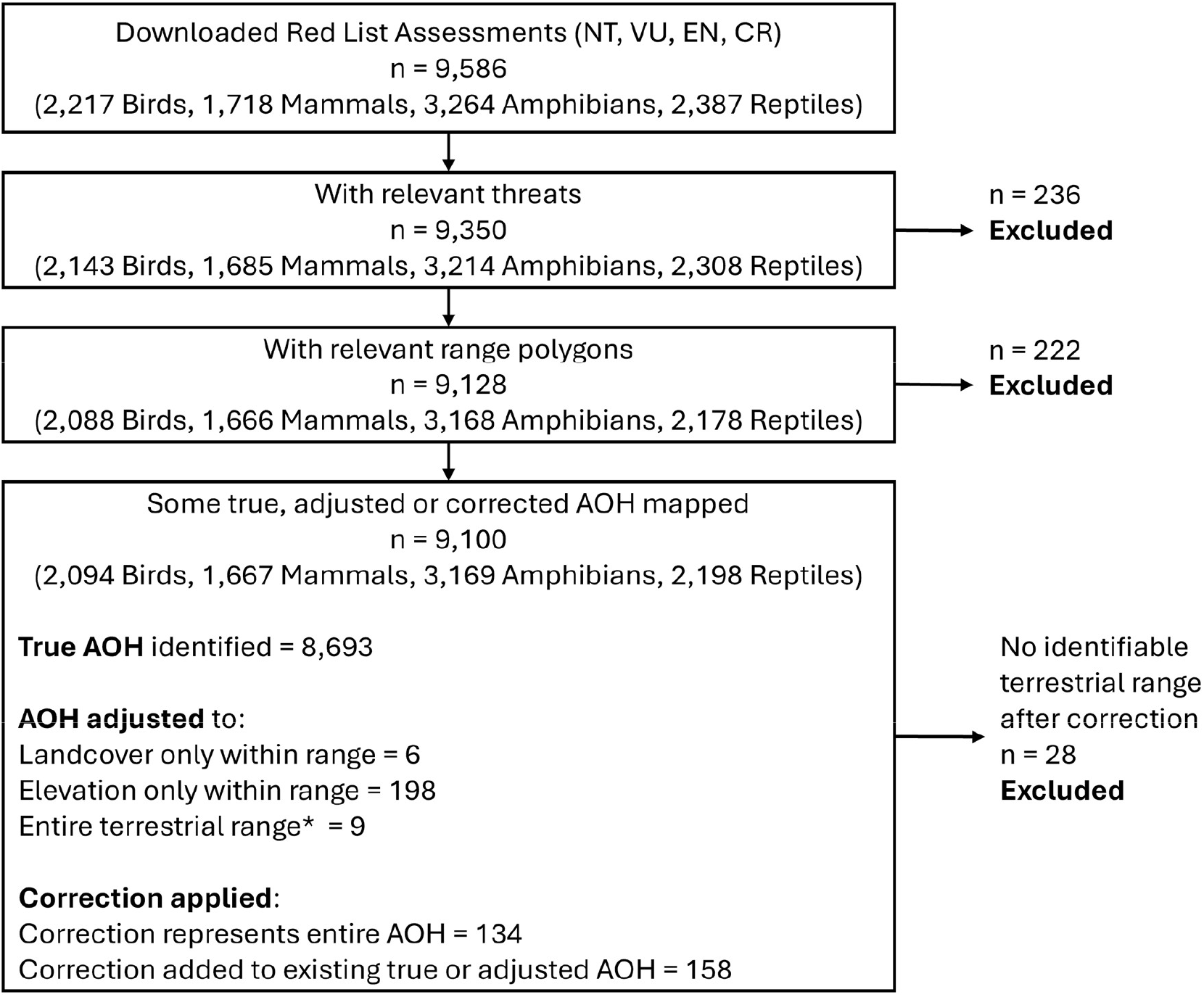
Summary of the Species included at each stage of data filtering and types of AOH maps included in the final calculation of STAR_T_.

A total of 515 species had a true AOH map and sufficient data to use in point validation (288 birds, 115 mammals, 57 amphibians and 55 reptiles). Mean point prevalence (proportion of species’ occurrences within the AOH) was 0.96 (SD 0.15) overall and 0.95 (SD 0.15) for birds, 0.97 (SD 0.11) for mammals, 0.96 (SD 0.14) for amphibians and 0.93 (SD 0.18) for reptiles.

A total of 505 species’ AOH maps (98% of those included in point validation) performed better than expected if AOH were distributed within the range at random. This left 10 that had a lower point prevalence than map prevalence (6 birds, 2 mammals, 2 amphibians and 0 reptiles; Fig. 2; also see Appendix S2).

**Figure 2.**
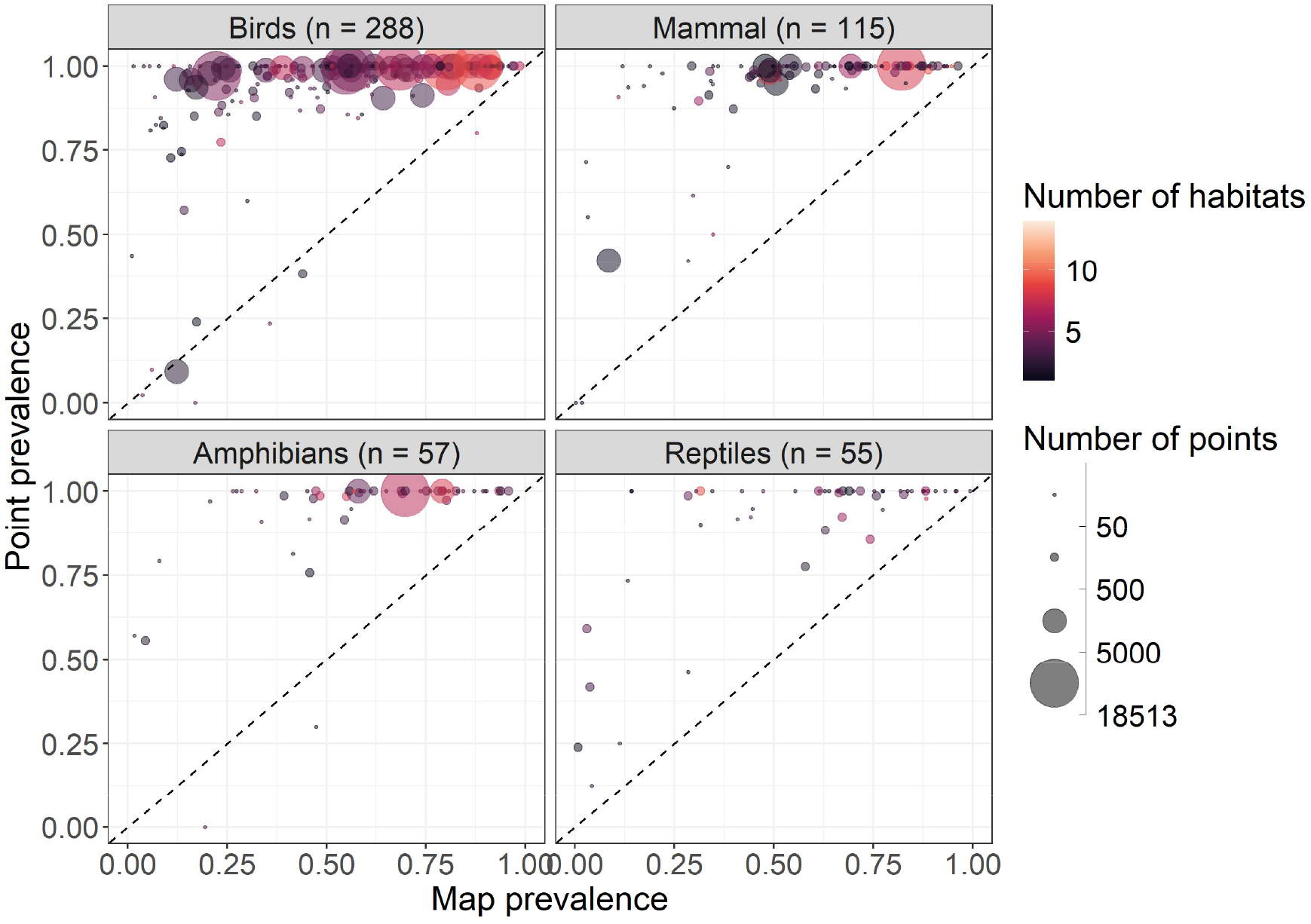
Validation of the area of habitat (AOH) mapping method for birds, mammals, amphibians and reptiles assessed as the relationship between point prevalence (the proportion of points where AOH was predicted to be) and map prevalence (the proportion of the species’ range that was predicted to be suitable). The dashed line indicates where map prevalence equals point prevalence and any AOH map above this line are considered to perform better than expected if AOH were distributed within the range at random.

### 4.2 Global threat abatement potential (STAR_T_) for terrestrial vertebrates

The total threat abatement potential (STAR_T_) score across 9,100 species (Table 3) was 2,204,100, of which 18.5% was contributed by birds, 17.9 % by mammals, 39.3 % by amphibians and 24.3% by reptiles.

**Table 3.** The number of species on which STAR_T_ was calculated per extinction risk category per taxonomic group.

| Extinction Risk | Birds | Mammals | Amphibians | Reptiles | Total |
| --- | --- | --- | --- | --- | --- |
| Near Threatened | 864 | 368 | 403 | 499 | 2,134 |
| Vulnerable | 655 | 542 | 789 | 558 | 2,544 |
| Endangered | 369 | 531 | 1,228 | 741 | 2,869 |
| Critically Endangered | 200 | 225 | 748 | 380 | 1,553 |
| **Total unique species** | **2,088** | **1,666** | **3,168** | **2,178** | **9,100** |

Threat abatement potential was geographically skewed (Fig. 3) with 12 countries making up 50% of the global threat abatement potential. These were: Brazil (6.3%), Madagascar (5.9%), Mexico (5.6%), Ecuador (5.5%), Indonesia (5.3%), Colombia (5.1%), India (4.3%), China (3.2%), Australia (2.9%), Sri Lanka (2.8%), Peru (2.7%) and Venezuela (Bolivarian Republic of) (2.6%) (Table 3). The lowest scoring 108 countries/territories together contributed only 1% of the total global threat abatement potential.

**Figure 3.**
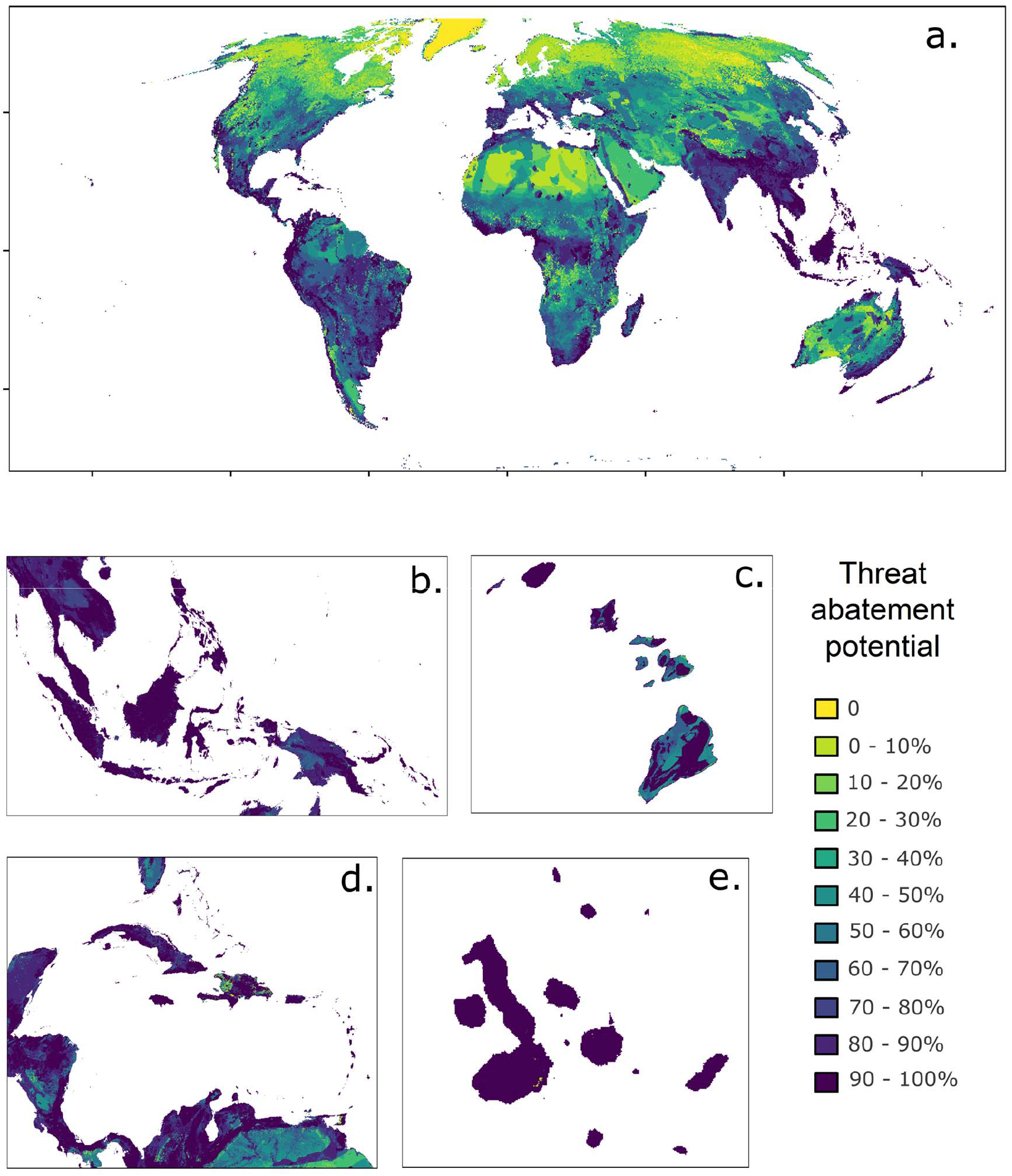
Total threat abatement potential (STAR_T_) for birds, mammals, amphibians and reptiles in each 1 km^2^ presented in Mollweide Equal Area Projection System a) globally, b) in South East Asia, c) in Hawaii, d) in the Caribbean, and e) in the Galapagos islands. Panels b-e represent a selection of areas with high STAR_T_ scores that are difficult to see when viewed globally. Aside from absolute 0, cells are coloured on a quantile scale where the lightest colour indicates the lowest 10% of non-zero scores and the darkest colour represents the top 10% of scores.

For 26 species, some current AOH was mapped but none intersected a GADM polygon with a ISO-3166 standard code where the species was present according to its Red List assessment. These species were included in calculating STAR_T_ but did not contribute to the score of any countries or territories. For example, the ranges of six of the 26 affected species were in Arunachal Pradesh; a disputed territory without an ISO-3166 standard code.

### 4.3 Which threats contribute the most to species extinction risk

The five highest ranking threats collectively contributed 64.2% of the global STAR_T_ score. These were: annual & perennial non-timber crops (21.2%), invasive & other problematic species, genes, or diseases (13.8%), logging & wood harvesting (12.8%), residential & commercial development (8.6%) and livestock farming & ranching (7.8%). Annual & perennial non-timber crops was the top threat in all major geographic regions, except for Oceania and Europe where the top threat was Invasive species, and Antarctica where the top threat was climate change (Fig. 4).

**Figure 4.**
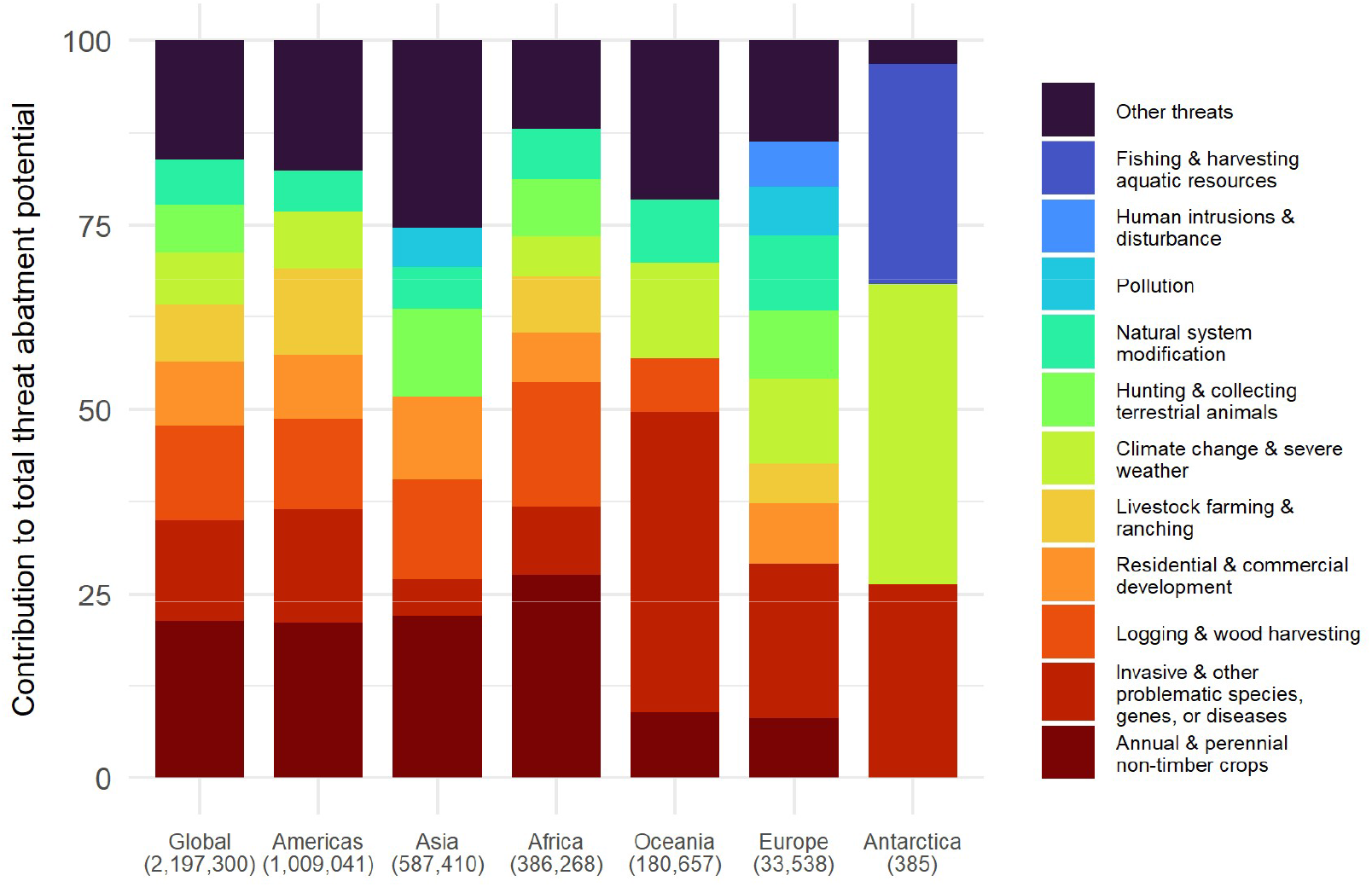
The contribution of different threats to the total threat abatement potential globally and for each UN major geographic region. Any threat that contributed less than 5% of the total score for the region was grouped with ‘Other threats’. Total STAR_T_ for each region is in parentheses. The global total here excludes the 26 species for which no AOH fell within any country or territory polygon, making it lower than the true global total.

### 4.4 Differences in STAR_T_ between original publication and now

The contribution of different threats to the global total was similar in STAR_T_v2 to STAR_T_v1, except that in STAR_T_v2 invasive species, ranked above logging, and residential & commercial development, ranked above livestock farming. The six countries that contributed the most to the global total threat abatement potential were the same in STAR_T_v2 as in STAR_T_v1, albeit in a different order (Table 4).

**Table 4.** Threat abatement scores in descending order by total STAR_T_ score for all countries that collectively contributed over 50% of the global threat abatement potential.

| ISO3 code | Country | STAR T v2 | % Global STAR T v2 | % Global STAR T v1 |
| --- | --- | --- | --- | --- |
| BRA | Brazil | 139,086.8 | 6.3 | 5.2 |
| MDG | Madagascar | 129,024.0 | 5.9 | 6.0 |
| MEX | Mexico | 123,425.0 | 5.6 | 6.1 |
| ECU | Ecuador | 120,315.6 | 5.5 | 4.1 |
| IDN | Indonesia | 115,949.0 | 5.3 | 7.1 |
| COL | Colombia | 111,196.0 | 5.1 | 7.0 |
| IND | India | 94,705.5 | 4.3 | 3.5 |
| CHN | China | 69,684.7 | 3.2 | 3.4 |
| AUS | Australia | 63,017.6 | 2.9 | 3.2 |
| LKA | Sri Lanka | 62,226.7 | 2.8 | 1.7 |
| PER | Peru | 59,440.4 | 2.7 | 4.1 |
| VEN | Venezuela (Bolivarian Republic of) | 57,034.6 | 2.6 | 2.2 |

Increases in threat abatement potential between STAR_T_v1 and STAR_T_v2 were observed in most 1 km^2^ cells (Fig. 5). Increases were observed in arid regions such as Northern Africa, Western Australia, and the Arabian Peninsula, as well as in densely forested regions such as the Congo basin. Notably widespread increases were observed over South and South-eastern Asia. Decreases were observed across much of Russia, Scandinavia, the Amazon, and an area of the Sahara Desert that covered substantial portions of Libya and Egypt. Though large increases and decreases were observed in some cells, the vast majority (97% of cells) observed changes between -0.81 and +0.95. It is important to note when comparing STAR_T_v2 to STAR_T_v1 that differences reflect both actual changes in threats or extinction risk, and changes in knowledge about these (see Discussion).

**Figure 5.**
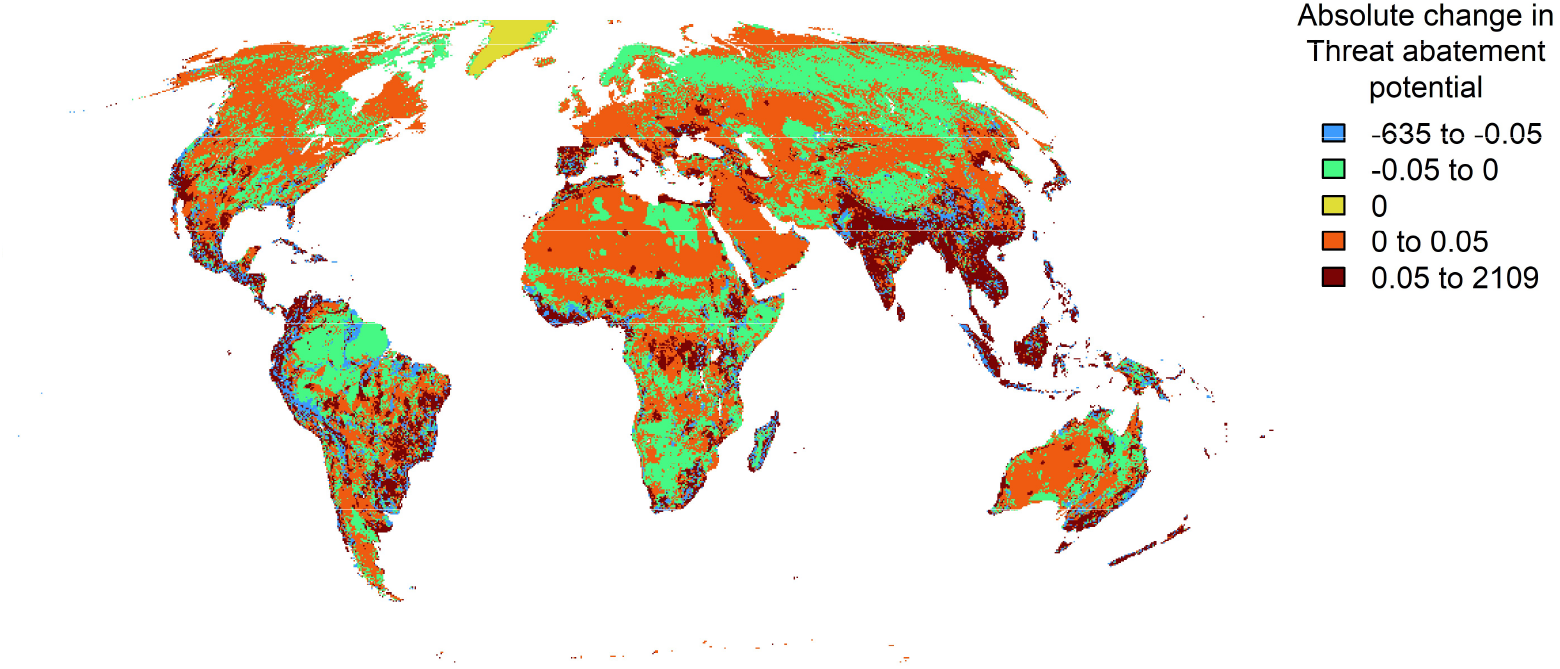
Change in threat abatement potential between STAR_T_v1 and STAR_T_v2 per 5 x 5 km cell presented in Mollweide equal area projection system.

## 5. Discussion

We present a substantially enhanced global estimate of threat abatement potential for terrestrial vertebrates based on recent advancements in data quality and data-driven methodologies, with expanded taxonomic scope (now including reptiles) and increased spatial resolution (now at 1 km^2^ resolution). The new data-driven method for mapping area of habitat (AOH), developed using data on birds and mammals (Lumbierres, Dahal, Di Marco, et al., 2022), performed similarly well when applied to amphibians and reptiles -the first time this has been demonstrated. We estimated that targeted actions to tackle a few high-impact threats in a few discrete locations, and cumulative efforts across multiple areas with lower individual potential, are both required to meaningfully reduce species’ extinction risk through threat abatement.

Changes in score from STAR_T_v1 to STAR_T_v2 represent enhanced understanding based on higher quality data and more data-driven methods, and do not necessarily indicate a change in threat intensity or extinction risk through time. First, the increases in threat abatement potential, particularly in many arid regions, are likely attributable to the inclusion of reptile species, which are more prevalent in drylands (Cox et al., 2022). Second, STAR_T_v2 includes data on 3,741 more species than STAR_T_v1, owing partially to reptiles (2,178 species), but also newly Red List assessed species, species that were previously assessed as Least Concern but are now classified as threatened or Near Threatened, and species for which relevant geographic range and AOH maps were not previously available. Second, some species included in STAR_T_v1 may not be included in STAR_T_v2 for the opposite reasons e.g. they have been downlisted to Least Concern. Third, some changes occur where taxonomic reclassification results in splitting or combining species, affecting their Red List assessments.

For these reasons, careful interpretation of the updated threat abatement scores is required. For example, STAR_T_v2 estimated that invasive species contributed a greater proportion of the global threat abatement potential than was estimated in STAR_T_v1. However, we cannot conclude from this there was an increase in the impact of invasive species on terrestrial vertebrates since 2019.

The Lumbierres, Dahal, Di Marco, et al. (2022) AOH mapping method performed similarly well on amphibians and reptiles as on birds and mammals, except that more amphibian species’ AOH maps were identified as outliers. There are several reasons why this might be the case. First, amphibian and reptile species tend to have smaller geographic range polygons than birds and mammals, amplifying errors among multiple sources of spatial data by making misalignment more likely and impactful. Second, amphibians are more reliant on microhabitats and microclimates, which are poorly represented by global land-cover datasets (Ficetola, Rondinini, Bonardi, Baisero, & Padoa-Schioppa, 2015). Third, the lower classification accuracy for mapping AOH for inland wetland habitat observed by Lumbierres, Dahal, Di Marco, et al. (2022) may disproportionately affect amphibian species. Finally, there were more amphibian species included than any other group making amphibians more likely among the outliers. Acknowledging that the availability of points was relatively low, particularly for reptiles and amphibians, our AOH maps performed slightly better than previous work, which found 77–94% of occurrences fell within the species’ AOH (point prevalence) and 79–95% of maps performed better than random (Dahal et al., 2022; Ficetola et al., 2015; Lumbierres, Dahal, Soria, et al., 2022; Rondinini et al., 2011).

The consideration of terrestrial systems in isolation (i.e. separately from marine and freshwater) introduces uncertainties around coastlines and supra-and inter-tidal habitats. The Lumbierres, Dahal, Di Marco, et al. (2022) crosswalk is restricted to terrestrial classes within the IUCN habitat classification scheme and excludes class 13 ‘Marine coastal/supratidal’, which encompasses habitats like sea cliffs and rocky offshore islands, caves, and sand dunes. This means that AOH for any species that also uses these habitats, such as cliff-breeding birds, will be an underestimate of the species’ true global AOH. Such habitats are also likely to be disproportionately affected by the error in missing CGLS100 terrestrial land-cover on islands. Building on the work to estimate STAR_T_ in the marine realm (Turner et al., 2024) and map AOH for inland wetland habitats (Ridley et al., 2025), future work should seek to generate a crosswalk that integrates habitats of all systems (terrestrial, marine and freshwater) to allow comparisons of threat abatement potential across realms.

Among an extensive suite of biodiversity-related metrics available (Burgess et al., 2024), STAR is identified as one of two that provide a necessary bottom-up approach to complement numerous top-down measures (Hawkins et al., 2024). The most similar alternative is the LIFE metric (Eyres et al., 2025). Conversely to STAR, LIFE includes Least Concern species and estimates extinction risk based on Durán et al. (2020) ‘s Global persistence score, in which extinction risk is estimated to non-linearly relate to the proportion of a species’ original AOH remaining (Eyres et al., 2025). It does not include other criteria of extinction risk embodied in IUCN Red List assessments such as geographic range, population size or rate of decline (IUCN, 2012b).

Estimated STAR_T_ provides essential data to assess potential to reduce extinction risk by tackling threats, thus supporting site selection, site-level calibration, target-setting and delivery of progress (Mair et al., 2026). Such estimates are useful to scope the potential across large land areas (or land portfolios) and large numbers of species. When followed by site-level calibration using locally collected data, site scale action can be related to site potential and global goals. This update has met multiple aspects of user-driven demand such as increased taxonomic scope, spatial resolution and data-driven methods. Such ongoing improvement is essential for uptake and facilitating the science-based reporting of contributions by cross-sectoral actors towards the KMGBF.

## Supporting information

Supplementary File 2 - Results and validation

## 6. Acknowledgements

We acknowledge the many individuals who have contributed their time and expertise to the Red List Assessments that were used in this work, in addition to those who supported the initial development of the STAR metric and related ongoing work. These individuals include but are not limited to Bernardo Strassburg, Bruce Young, Craig Hilton-Taylor, Caroline Pollock and Richard Jenkins. We would like to acknowledge the generous funding provided by the Global Environment Facility (GEF) through the Knowledge-4-Nature project (10897) that has supported the production of this work and FAR. LM was funded by Newcastle University.

## 7. Data availability statement

The data and code to accompany this research are available in the online repository (https://zenodo.org/records/17989947). This includes: species AOH maps at 1 km resolution (.tif), details of species filtering and validation results for AOH maps generated (.csv), global maps of threat abatement potential in total and for each threat at 1km resolution in Mollweide Equal Area Projection system (.tif), a table of the STAR_T_ score per species per threat per gadm country, and R scripts to reproduce the full workflow including calculating area of habitat and STAR_T_. For commercial purposes this data must be accessed via the Integrated Biodiversity Assessment Tool (IBAT, www.ibat-alliance.org), as with any IUCN Red List-derived data.

## 8. Author contributions

Conceptualisation: FAR, LM, TB, NM, TB, SB, FH, LB, PM, TS, JT

Data curation: JC, KN, NC, CR, MP, VM, JPR

Formal Analysis, investigation & validation: FAR, MWD, RJ

Funding Acquisition & project administration: TB, NM, AR, RJ

Methodology: FAR, LM, TS, JT, DB, SB, RJ, MWD, ST, NM

Project Administration & software: NM, AR, RJ, ST

Writing – original draft preparation: FAR

Writing – review & editing: All authors reviewed, edited and approved the final manuscript

