## Supplementary File 1 - Methods and Tables for "Global threat abatement potential for terrestrial vertebrates"

### Appendix 2. Supplementary results and validation

#### Adjustments and corrections to species' AOH maps

The AOH mapping method was applied to 9,128 species, including 2,094 birds, 1,667 mammals, 3,169 amphibians and 2,198 reptiles. This includes the 28 species for which no terrestrial land-cover could be located within the range and were excluded from the calculation of $STAR_T$ (Table S2.1).

Adjustments to the definition of AOH and corrections due to errors in the land-cover data were applied to a total of 564 species (Table S2.2). These totals include 65 species for which there was no suitable elevation within the range but there was some suitable land-cover, so AOH was adjusted to represent suitable land-cover only within the species' terrestrial range. For 198 species, suitable elevation was identified within the species' terrestrial range but no suitable land-cover, so AOH was adjusted to represent suitable elevation only within the species' terrestrial range. For 172 species, the whole of the species' terrestrial range was considered to contain potentially suitable habitat because a) areas at suitable elevation did not coincide with those containing suitable land-cover (7 species), b) neither suitable land-cover nor suitable elevation was identified within the species' terrestrial range (2 species) or c) no cells within the species' range were at least 50% covered by any CGLS100 terrestrial land-cover classes (162 species) and on which the correction was applied. This includes the 28 species for which the correction was unsuccessful in identifying any terrestrial area within the species' range. A further, 158 species were found to be affected by the error in missing CGLS100 terrestrial land-cover but for which some AOH (true AOH or adjusted) was already mapped elsewhere.

For 26 species, some current AOH was mapped but none intersected a country or territory that corresponded to the countries attributed in the IUCN Red List Assessment for the species (Table S2.3). These species were included in calculating $STAR_T$ but did not contribute to the score of any countries or territories.

#### Potential errors in species' elevation limits

An elevation adjustment was applied to 445 species in total (see main text). Of these, some suitable elevation was identified for407 species (91.46%). No suitable elevation was identified within the range of 67 species, of which 29 species (43.28%) had a raw elevation range less than 12m and their elevation limits were adjusted. A total of 369 species had an elevation range of zero in the raw data. It was concluded that the error was due to the elevation coding and this information was fed back to the relevant IUCN Red List authority.

**Table S2.1.** Details of the 28 species for which no terrestrial land-cover could be located within the species' range even after adjustments and corrections were applied. Current AOH was not identifiable for these species and as such they were excluded from $STAR_T$.

| TaxonID | Scientific name | Taxonomic group |
|---|---|---|
| 939 | Alsophis antiguae | REPTILIA |
| 74995219 | Anolis ernestwilliamsi | REPTILIA |
| 74995581 | Anolis kahouannensis | REPTILIA |
| 14811 | Carinascincus palfreymani | REPTILIA |
| 136520 | Cavia intermedia | MAMMALIA |
| 101742350 | Cyrtodactylus jarakensis | REPTILIA |
| 203507 | Enulius bifoveatus | REPTILIA |
| 12080 | Erythrolamprus ornatus | REPTILIA |
| 103815245 | Geospiza septentrionalis | AVES |
| 22692441 | Hypotaenidia owstoni | AVES |
| 196581 | Lepidodactylus oligoporus | REPTILIA |
| 196582 | Lepidodactylus paurolepis | REPTILIA |
| 196585 | Lepidodactylus tepukapili | REPTILIA |
| 47102814 | Mabuya cochonae | REPTILIA |
| 22711063 | Mimus trifasciatus | AVES |
| 17424269 | Nactus serpensinsula | REPTILIA |
| 156730274 | Oligosoma salmo | REPTILIA |
| 120190169 | Oligosoma taumakae | REPTILIA |
| 136140 | Ololygon faivovichi | AMPHIBIA |
| 174139 | Pholidoscelis corvinus | REPTILIA |
| 8218 | Plestiodon longirostris | REPTILIA |
| 157290 | Podarcis levendis | REPTILIA |
| 22734320 | Pomarea mendozae | AVES |
| 22707192 | Pomarea whitneyi | AVES |
| 47103206 | Spondylurus macleani | REPTILIA |
| 47103260 | Spondylurus monitae | REPTILIA |
| 22720722 | Telespiza ultima | AVES |
| 110230530 | Trachylepis adamastor | REPTILIA |

**Table S2.2.** The number of species for which each adjustment and/or correction to AOH was applied. Adjustments refer to changes in what the AOH represented where no true AOH (suitable land-cover at suitable elevation within the range) was located whereas, corrections attempted to overcome errors of missing terrestrial land-cover in the Copernicus (100m) Global Land-cover data.

| Adjustment | Correction | Number of species |
|---|---|---|
| None | None | 8,564 |
| AOH adjusted to elevation within range | None | 171 |
| AOH adjusted to range | Existing range and AOH corrected to the area of intersection between the range and affected GADM polygons | 134 |
| None | Existing range and AOH corrected to additionally include the area of intersection between the range and affected GADM polygons | 129 |
| AOH adjusted to landcover within range | None | 64 |
| AOH adjusted to range | No affected GADM polygons intersected the range | 28 |
| AOH adjusted to elevation within range | Existing range and AOH corrected to additionally include the area of intersection between the range and affected GADM polygons | 27 |
| AOH adjusted to range | None | 9 |
| AOH adjusted to landcover within range | Existing range and AOH corrected to additionally include the area of intersection between the range and affected GADM polygons | 1 |
| AOH adjusted to range | Existing range and AOH corrected to additionally include the area of intersection between the range and affected GADM polygons | 1 |

**Table S2.3.** Details of the species for which some AOH was mapped but not intersected any country boundary according to the GADM polygon dataset.

| TaxonID | Scientific name | AOH (Number of cells) | Map prevalence | AOH Adjustments & Corrections | Manual investigation notes |
|---|---|---|---|---|---|
| 165414242 | Nanorana arunachalensis | 3,837.71 | 0.78 | | Range falls within Arunachal Pradesh - possible disputed territory |
| 150087712 | Megophrys himalayana | 22.04 | 0.82 | | Range falls within Arunachal Pradesh - possible disputed territory |
| 150087897 | Megophrys periosa | 665.83 | 0.93 | | Range falls within Arunachal Pradesh - possible disputed territory |
| 88368304 | Megophrys vegrandis | 32.89 | 0.52 | | Range falls within Arunachal Pradesh - possible disputed territory |
| 22680203 | Anas laysanensis | 0.69 | 0.17 | | Small outlying island of Hawaii archipelago - range map seems accurately positioned relative to land in basemap on Red List website - possible mismatch with gadm polygon |

| TaxonID | Scientific name | AOH (Number of cells) | Map prevalence | AOH Adjustments & Corrections | Manual investigation notes |
|---|---|---|---|---|---|
| 22714797 | Acrocephalus familiaris | 0.69 | 0.17 | | Small outlying island of Hawaii archipelago - possible mismatch with gadm polygon |
| 22696876 | Leucocarbo ranfurlyi | 1.00 | 1.00 | AOH reverted to elevation within range | Small outlying island of New Zealand - possible spatial mismatch with range and gadm polygon |
| 22734628 | Liocichla bugunorum | 264.28 | 0.77 | | Range falls within Arunachal Pradesh - possible disputed territory |
| 103798418 | Poodytes caudatus | 0.82 | 0.82 | | Small outlying island of New Zealand - possible spatial mismatch with range and gadm polygon |
| 103734886 | Petroica dannefaerdi | 1.00 | 1.00 | AOH reverted to elevation within range | Small outlying island of New Zealand - possible spatial mismatch with range and gadm polygon |
| 22682531 | Rhyticeros narcondami | 6.75 | 0.84 | | Narcondam island - potential range map positional inaccuracy and/or gadm polygon |
| 22720728 | Telespiza cantans | 2.22 | 0.56 | | Papahanaumokuakea Marine Nat'l Mon - potential spatial inaccuracy in gadm polygon |
| 14563 | Mogera uchidai | 4.00 | 1.00 | AOH reverted to elevation within range | Small island off Japan - potential inaccuracy in range map position and gadm polygon |
| 45959013 | Petaurista mechukaensis | 7,453.13 | 0.58 | | Range falls within Arunachal Pradesh - possible disputed territory |
| 16657 | Peromyscus dickeyi | 3.41 | 0.49 | | Isla Tortuga - potential inaccuracy in gadm polygon |
| 178336 | Anolis nubilus | 0.07 | 0.07 | | Also Redonda - range map position seems accurate relative to basemap on Red List website - supports spatial mismatch with gadm |
| 74994414 | Anolis altavelensis | 1.00 | 1.00 | AOH reverted to elevation within range | Small outlying island of Dominican Republic - possible spatial inaccuracies in range and/or gadm polygon |
| 44577361 | Anolis gorgonae | 0.47 | 0.47 | | Small offshore island of colombia - range map seems accurately positioned relative to land in basemap on Red List website - possible mismatch with gadm polygon |
| 104682048 | Cnemaspis boulengerii | 21.06 | 0.27 | | Offshore island potential inaccuracy in gadm polygon |

| TaxonID | Scientific name | AOH (Number of cells) | Map prevalence | AOH Adjustments & Corrections | Manual investigation notes |
|---|---|---|---|---|---|
| 104680418 | Dibamus kondaoensis | 20.70 | 0.29 | | Offshore island potential inaccuracy in gadm polygon and in range polygon |
| 75306189 | Leiocephalus altavelensis | 1.00 | 1.00 | AOH reverted to elevation within range | Small outlying island of Dominican Republic - possible spatial inaccuracies in range and/or gadm polygon |
| 102345892 | Eutropis quadratilobus | 0.10 | 0.01 | | Extremely small range, buffer around point, not positioned in Bhutan |
| 44579522 | Pseudogonatodes peruvianus | 153.70 | 0.13 | | Range falls in Peru |
| 50009685 | Pholidoscelis atratus | 0.07 | 0.07 | | Small outlying island Antigua and Barbuda (Redonda) - potential mismatch in position of range, position of land and/or position of gadm polygon |
| 120190180 | Oligosoma tekakahu | 13.00 | 1.00 | AOH reverted to elevation within range | Offshore island potential inaccuracy in gadm polygon |
| 75606372 | Tropidophis bucculentus | 4.00 | 1.00 | AOH reverted to elevation within range | Small island between Jamaica, Haiti and Cuba - possible inaccuracy of country coding or gadm polygon |

#### Map prevalence and identification of outliers

Among the 8,693 species for which some suitable habitat at suitable elevation was identified within the range (i.e true AOH), the mean map prevalence was 0.61 (SD 0.27). This varied among taxonomic groups. The mean map prevalence was 0.6 (SD 0.27) for birds, 0.65 (SD 0.26) for mammals, 0.64 (SD 0.26) for amphibians and 0.52 (SD 0.29) for reptiles. The difference between observed values for map prevalence and those predicted by the logistic regression model identified 16 species to be outliers (1 bird, 2 mammals, 13 amphibians and 0 reptiles) (Table S2.4). A further 131 species had very low map prevalence (< 1% of inland range was AOH) (Table S2.5).

**Table S2.4.** Species identified as outliers based on the difference between observed and predicted map prevalence

| TaxonID | Scientific name | Class | Range size (Number of cells) | Map prevalence | Elevation range (m) | Habitats |
|---|---|---|---|---|---|---|
| 977 | Alytes muletensis | AMPHIBIA | 182 | 0.01 | 840 | 14.2: Pasture, 15: Artificial Aquatic, 5: Inland Wetlands |
| 41032 | Litoria cooloolensis | AMPHIBIA | 2,300 | 0.02 | 8,580 | 5: Inland Wetlands, Other |
| 41143 | Crinia tasmaniensis | AMPHIBIA | 23,857 | 0.05 | 1,160 | 15: Artificial Aquatic, 5: Inland Wetlands |

| TaxonID | Scientific name | Class | Range size (Number of cells) | Map prevalence | Elevation range (m) | Habitats |
|---|---|---|---|---|---|---|
| 58514 | Ptychadena newtoni | AMPHIBIA | 325 | 0.01 | 600 | 14.1: Arable, 14.3: Plantations, 14.4: Rural Gardens, 14.5: Urban, 14.6: Heavily Degraded Former Forest, 15: Artificial Aquatic, 5: Inland Wetlands |
| 58567 | Pelophylax cerigensis | AMPHIBIA | 9 | 0.01 | 500 | 15: Artificial Aquatic, 5: Inland Wetlands |
| 58873 | Philautus microdiscus | AMPHIBIA | 14 | 0.02 | 122 | 1: Forest |
| 59283 | Gyrinophilus subterraneus | AMPHIBIA | 7 | 0.07 | 9,007 | 5: Inland Wetlands, Other |
| 59284 | Eurycea wallacei | AMPHIBIA | 248 | 0.05 | 40 | 5: Inland Wetlands, Other |
| 59391 | Aquiloeurycea praecellens | AMPHIBIA | 19 | 0.13 | 9,007 | 1: Forest |
| 136582 | Habromys ixtlani | MAMMALIA | 125 | 0.97 | 500 | 1: Forest |
| 164778 | Bufotes zugmayeri | AMPHIBIA | 29,149 | 0.06 | 9,007 | 14.1: Arable, 5: Inland Wetlands |
| 22698097 | Pterodroma caribbaea | AVES | 4 | 1.00 | 7,580 | 1: Forest, Other, Other |
| 48100899 | Rhacophorus spelaeus | AMPHIBIA | 5,392 | 0.10 | 8,440 | 5: Inland Wetlands, Other |
| 49583046 | Stumpffia be | AMPHIBIA | 323 | 0.01 | 8,490 | 5: Inland Wetlands, Other |
| 112139816 | Rattus detentus | MAMMALIA | 1,900 | 0.00 | 166 | 14.3: Plantations, 14.4: Rural Gardens, 14.6: Heavily Degraded Former Forest |
| 178696039 | Pristimantis torresi | AMPHIBIA | 418 | 0.04 | 2,233 | 3: Shrubland |

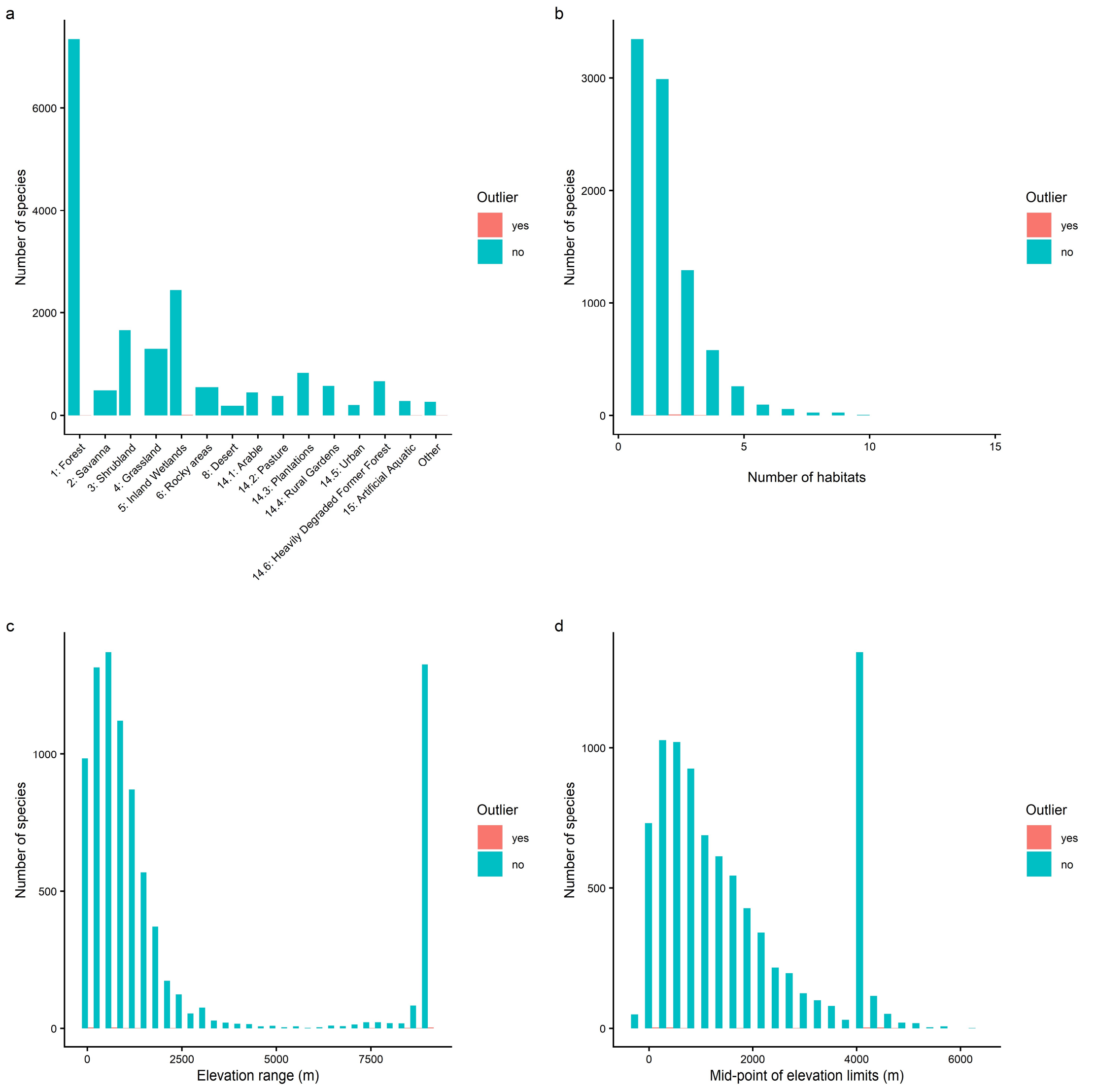


**Figure S2.1.** Comparison between species identified as outliers and species not identified as outliers based on the difference between observed and predicted map prevalence a) Habitats, b) number of habitats, c) elevation range, d) elevation mid-point.

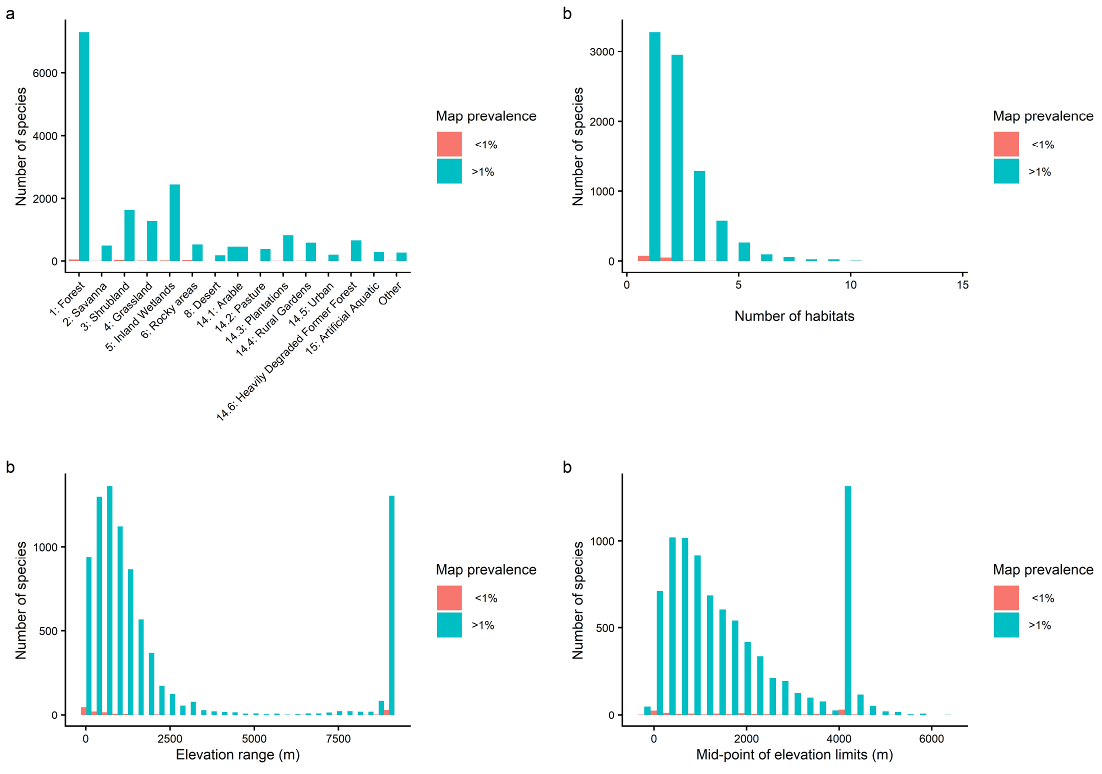


**Figure S2.2** Comparison between species where true AOH covered less than 1% of the range and those where AOH covered more than 1%.

**Table S2.5.** Species for which some true AOH was mapped but it covered less than 1% of the range

| TaxonID | Scientific name | Class | Range size (Number of cells) | Map prevalence | Elevation range (m) | Habitats |
|---|---|---|---|---|---|---|
| 4420 | Chalinolobus neocaledonicus | MAMMALIA | 16,689 | 0.0062 | 8,580 | 14.4: Rural Gardens, 14.5: Urban, 3: Shrubland |
| 5588 | Crocidura zimmermanni | MAMMALIA | 1,224 | 0.0062 | 1,260 | 6: Rocky areas, Other |
| 5862 | Ctenotus zastictus | REPTILIA | 74 | 0.0077 | 9,007 | 4: Grassland |
| 6287 | Dasyprocta ruatanica | MAMMALIA | 156 | 0.0092 | 9,007 | 1: Forest |
| 6607 | Dinaromys bogdanovi | MAMMALIA | 80,719 | 0.0016 | 2,200 | 6: Rocky areas |
| 8766 | Furcifer minor | REPTILIA | 4,018 | 0.0002 | 590 | 14.3: Plantations, 1: Forest |
| 12005 | Liolaemus lorenzmuelleri | REPTILIA | 1,600 | 0.0090 | 300 | 4: Grassland |
| 12267 | Lonchophylla mordax | MAMMALIA | 21,456 | 0.0094 | 9,007 | 1: Forest, Other |
| 12271 | Lonchorhina fernandezi | MAMMALIA | 108,737 | 0.0004 | 20 | 2: Savanna, Other |
| 12272 | Lonchorhina marinkellei | MAMMALIA | 166,838 | 0.0009 | 60 | 4: Grassland, Other |
| 14189 | Myotis peninsularis | MAMMALIA | 3,345 | 0.0077 | 2,200 | 1: Forest, Other, Other |
| 16750 | Petrogale xanthopus | MAMMALIA | 26,597 | 0.0013 | 9,007 | 6: Rocky areas |
| 16753 | Petrogale sharmani | MAMMALIA | 17,654 | 0.0003 | 9,007 | 6: Rocky areas |

| TaxonID | Scientific name | Class | Range size (Number of cells) | Map prevalence | Elevation range (m) | Habitats |
|---|---|---|---|---|---|---|
| 18113 | Praomys hartwigi | MAMMALIA | 8,246 | 0.0039 | 200 | 1: Forest |
| 20485 | Callospermophilus madrensis | MAMMALIA | 61,966 | 0.0096 | 750 | 1: Forest |
| 22998 | Montivipera wagneri | REPTILIA | 2,963 | 0.0022 | 800 | 6: Rocky areas |
| 23000 | Vipera darevskii | REPTILIA | 207 | 0.0006 | 600 | 14.2: Pasture, 6: Rocky areas |
| 29454 | Phasmasaurus maruia | REPTILIA | 872 | 0.0010 | 840 | 3: Shrubland, 6: Rocky areas |
| 41310 | Sylvilagus robustus | MAMMALIA | 39,505 | 0.0052 | 847 | 1: Forest |
| 41514 | Petrogale godmani | MAMMALIA | 37,399 | 0.0010 | 9,007 | 6: Rocky areas |
| 54553 | Atelopus sonsonensis | AMPHIBIA | 414 | 0.0044 | 12 | 1: Forest, 5: Inland Wetlands |
| 54568 | Rhinella amabilis | AMPHIBIA | 12 | 0.0017 | 150 | 15: Artificial Aquatic, 5: Inland Wetlands |
| 54791 | Rhinella vellardi | AMPHIBIA | 335 | 0.0016 | 12 | 1: Forest, 5: Inland Wetlands |
| 55066 | Allobates chalcopis | AMPHIBIA | 18 | 0.0056 | 600 | 4: Grassland, 5: Inland Wetlands |
| 55260 | Aromobates serranus | AMPHIBIA | 106 | 0.0009 | 500 | 1: Forest, 5: Inland Wetlands |
| 56027 | Stefania ginesi | AMPHIBIA | 952 | 0.0060 | 750 | 5: Inland Wetlands, 6: Rocky areas |
| 56028 | Stefania goini | AMPHIBIA | 43 | 0.0033 | 300 | 3: Shrubland, 5: Inland Wetlands |
| 56037 | Stefania schuberti | AMPHIBIA | 627 | 0.0087 | 650 | 5: Inland Wetlands, 6: Rocky areas |
| 56220 | Hyperolius viridigulosus | AMPHIBIA | 32,378 | 0.0011 | 500 | 1: Forest, 5: Inland Wetlands |
| 56382 | Dischidodactylus colonnelloi | AMPHIBIA | 129 | 0.0033 | 12 | 3: Shrubland |
| 56730 | Eleutherodactylus lucioi | AMPHIBIA | 12 | 0.0025 | 12 | 1: Forest, 6: Rocky areas |
| 56909 | Eleutherodactylus rhodesi | AMPHIBIA | 123 | 0.0071 | 12 | 1: Forest |
| 57342 | Telmatobius huayra | AMPHIBIA | 17,468 | 0.0000 | 12 | 5: Inland Wetlands |
| 57354 | Telmatobius philippii | AMPHIBIA | 184 | 0.0033 | 100 | 4: Grassland, 5: Inland Wetlands |
| 58567 | Pelophylax cerigensis | AMPHIBIA | 9 | 0.0067 | 500 | 15: Artificial Aquatic, 5: Inland Wetlands |
| 58751 | Odorrana wuchuanensis | AMPHIBIA | 72 | 0.0075 | 12 | 5: Inland Wetlands, Other |
| 59050 | Ambystoma amblycephalum | AMPHIBIA | 33 | 0.0091 | 12 | 15: Artificial Aquatic, 1: Forest, 4: Grassland, 5: Inland Wetlands |
| 61813 | Pristimantis marahuaka | AMPHIBIA | 153 | 0.0018 | 12 | 3: Shrubland |
| 61820 | Pristimantis telefericus | AMPHIBIA | 36 | 0.0044 | 600 | 3: Shrubland |
| 64105 | Sceloporus goldmani | REPTILIA | 224 | 0.0021 | 9,007 | 4: Grassland |
| 64169 | Urosaurus clarionensis | REPTILIA | 26 | 0.0023 | 9,007 | 1: Forest |
| 64255 | Aspidoscelis catalinensis | REPTILIA | 32 | 0.0016 | 9,007 | 3: Shrubland, 6: Rocky areas, Other |
| 64314 | Crotalus catalinensis | REPTILIA | 32 | 0.0016 | 9,007 | 6: Rocky areas, 8: Desert |

| TaxonID | Scientific name | Class | Range size (Number of cells) | Map prevalence | Elevation range (m) | Habitats |
|---|---|---|---|---|---|---|
| 135741 | Microkayla katantika | AMPHIBIA | 22 | 0.0068 | 400 | 1: Forest, 3: Shrubland |
| 135828 | Eleutherodactylus juanariveroi | AMPHIBIA | 11 | 0.0009 | 12 | 5: Inland Wetlands |
| 136053 | Microkayla ankohuma | AMPHIBIA | 56 | 0.0062 | 150 | 3: Shrubland |
| 136101 | Microkayla harveyi | AMPHIBIA | 13 | 0.0015 | 600 | 3: Shrubland |
| 136145 | Microkayla kallawaya | AMPHIBIA | 22 | 0.0082 | 12 | 1: Forest, 3: Shrubland |
| 136165 | Microkayla chacaltaya | AMPHIBIA | 115 | 0.0047 | 300 | 3: Shrubland |
| 136463 | Petrogale purpureicollis | MAMMALIA | 82,986 | 0.0002 | 9,007 | 6: Rocky areas |
| 136509 | Petrogale mareeba | MAMMALIA | 20,729 | 0.0001 | 1,000 | 6: Rocky areas |
| 136638 | Heteromys oasicus | MAMMALIA | 2,627 | 0.0067 | 9,007 | 1: Forest |
| 136648 | Alticola olchonensis | MAMMALIA | 1,521 | 0.0026 | 9,007 | 6: Rocky areas |
| 136677 | Lepilemur petteri | MAMMALIA | 14,379 | 0.0037 | 23 | 1: Forest |
| 157271 | Iberolacerta galani | REPTILIA | 1,004 | 0.0002 | 900 | 6: Rocky areas |
| 164578 | Darevskia unisexualis | REPTILIA | 16,060 | 0.0049 | 2,427 | 6: Rocky areas |
| 164647 | Phrynocephalus persicus | REPTILIA | 131,606 | 0.0076 | 1,577 | 3: Shrubland, 4: Grassland, 8: Desert |
| 169686 | Rabdion forsteni | REPTILIA | 9,349 | 0.0010 | 12 | 1: Forest |
| 169880 | Gekko gigante | REPTILIA | 12 | 0.0017 | 200 | 6: Rocky areas, Other |
| 172750 | Paracontias minimus | REPTILIA | 42 | 0.0081 | 90 | 3: Shrubland |
| 172760 | Calumma tsaratananense | REPTILIA | 1,561 | 0.0010 | 350 | 3: Shrubland |
| 172788 | Trachylepis nancycoutuae | REPTILIA | 7,043 | 0.0086 | 50 | 3: Shrubland, 6: Rocky areas |
| 172998 | Grandidierina petiti | REPTILIA | 1,635 | 0.0035 | 20 | 1: Forest |
| 174131 | Eutropis ashwamedhi | REPTILIA | 927 | 0.0036 | 70 | 3: Shrubland |
| 177486 | Micrurus ruatanus | REPTILIA | 129 | 0.0002 | 20 | 1: Forest |
| 178408 | Contomastix vittata | REPTILIA | 11,303 | 0.0069 | 1,329 | 14.2: Pasture, 1: Forest |
| 178502 | Cryptoblepharus gloriosus | REPTILIA | 580 | 0.0021 | 12 | 3: Shrubland, Other |
| 178744 | Liolaemus arambarensis | REPTILIA | 721 | 0.0033 | 9,007 | 3: Shrubland, Other |
| 186104 | Bolitoglossa pygmaea | AMPHIBIA | 37 | 0.0003 | 100 | 4: Grassland |
| 192040 | Elaphe moellendorffi | REPTILIA | 332,744 | 0.0010 | 270 | 3: Shrubland, 6: Rocky areas |
| 194109 | Hemidactylus gujaratensis | REPTILIA | 3,215 | 0.0068 | 12 | 1: Forest, 6: Rocky areas |
| 203150 | Pristidactylus achalensis | REPTILIA | 7,044 | 0.0009 | 545 | 2: Savanna |
| 203305 | Oxybelis wilsoni | REPTILIA | 145 | 0.0020 | 95 | 1: Forest, 3: Shrubland |
| 203508 | Enulius roatanensis | REPTILIA | 37 | 0.0003 | 12 | 1: Forest |
| 15157465 | Apostolepis striata | REPTILIA | 32 | 0.0037 | 9,007 | 2: Savanna |

| TaxonID | Scientific name | Class | Range size (Number of cells) | Map prevalence | Elevation range (m) | Habitats |
|---|---|---|---|---|---|---|
| 16391176 | Marisora roatanae | REPTILIA | 17 | 0.0006 | 9,007 | 1: Forest |
| 18490428 | Hemidactylus greeffii | REPTILIA | 859 | 0.0002 | 1,400 | 14.5: Urban, Other |
| 22698252 | Puffinus huttoni | AVES | 32,938 | 0.0006 | 600 | 3: Shrubland, 4: Grassland, Other, Other |
| 22698530 | Hydrobates macrodactylus | AVES | 456 | 0.0000 | 9,007 | 1: Forest, Other, Other |
| 22700773 | Zaratornis stresemanni | AVES | 64,896 | 0.0007 | 2,060 | 1: Forest |
| 22701605 | Formicivora erythronotos | AVES | 172 | 0.0026 | 477 | 14.3: Plantations, 14.6: Heavily Degraded Former Forest, 3: Shrubland |
| 22703783 | Amytornis woodwardi | AVES | 5,882 | 0.0044 | 300 | 4: Grassland, 6: Rocky areas |
| 22716864 | Curruca buryi | AVES | 64,804 | 0.0001 | 1,400 | 1: Forest |
| 22723786 | Geospiza heliobates | AVES | 14 | 0.0007 | 8,580 | 1: Forest |
| 22727509 | Coenocorypha huegeli | AVES | 19 | 0.0053 | 250 | 3: Shrubland, 4: Grassland |
| 22736183 | Zosterops somadikartai | AVES | 386 | 0.0025 | 527 | 14.3: Plantations, 14.4: Rural Gardens, 14.6: Heavily Degraded Former Forest, 3: Shrubland |
| 42685911 | Brookesia brunoi | REPTILIA | 20 | 0.0005 | 12 | 1: Forest |
| 42760615 | Trioceros kinangopensis | REPTILIA | 34 | 0.0024 | 100 | 3: Shrubland |
| 44579029 | Enyalioides touzeti | REPTILIA | 1,689 | 0.0056 | 300 | 1: Forest |
| 47102633 | Brasiliscincus caissara | REPTILIA | 3,335 | 0.0018 | 9,007 | 3: Shrubland |
| 47103187 | Spondylurus lineolatus | REPTILIA | 368 | 0.0074 | 310 | 1: Forest |
| 47755599 | Alsophylax szczerbaki | REPTILIA | 17,007 | 0.0056 | 200 | 8: Desert |
| 47755615 | Alsophylax tadjikiensis | REPTILIA | 14,554 | 0.0068 | 100 | 3: Shrubland |
| 47992992 | Lophuromys pseudosikapusi | MAMMALIA | 1,936 | 0.0055 | 12 | 1: Forest |
| 48269619 | Cryptotis griseoventris | MAMMALIA | 192 | 0.0034 | 6,480 | 1: Forest |
| 48442737 | Phyllodactylus delsolari | REPTILIA | 4,131 | 0.0038 | 980 | 1: Forest |
| 48443921 | Phyllopezus maranjonensis | REPTILIA | 2,064 | 0.0078 | 100 | 1: Forest, 6: Rocky areas |
| 49583046 | Stumpffia be | AMPHIBIA | 323 | 0.0072 | 8,490 | 5: Inland Wetlands, Other |
| 49845682 | Tropidurus xanthochilus | REPTILIA | 1,072 | 0.0037 | 15 | 14.2: Pasture, 2: Savanna |
| 67610740 | Atractus multidentatus | REPTILIA | 28 | 0.0011 | 12 | 1: Forest |
| 74994592 | Anolis breslini | REPTILIA | 1,002 | 0.0086 | 9,007 | 1: Forest |
| 75091532 | Anolis garridoi | REPTILIA | 250 | 0.0016 | 100 | 2: Savanna |
| 75158906 | Amphisbaena arda | REPTILIA | 6,294 | 0.0005 | 9,007 | 1: Forest |

| TaxonID | Scientific name | Class | Range size (Number of cells) | Map prevalence | Elevation range (m) | Habitats |
|---|---|---|---|---|---|---|
| 75317710 | Leiocephalus rhutidira | REPTILIA | 58 | 0.0028 | 9,007 | 1: Forest |
| 75323721 | Leiocephalus vinculum | REPTILIA | 705 | 0.0078 | 591 | 1: Forest |
| 75605218 | Phyllodactylus sommeri | REPTILIA | 67 | 0.0049 | 310 | 1: Forest, 6: Rocky areas |
| 75605330 | Sphaerodactylus asterulus | REPTILIA | 254 | 0.0027 | 235 | 1: Forest |
| 75605475 | Sphaerodactylus lazelli | REPTILIA | 20 | 0.0085 | 12 | 1: Forest |
| 75605902 | Sphaerodactylus shrevei | REPTILIA | 479 | 0.0023 | 50 | 1: Forest, 6: Rocky areas |
| 77639796 | Adenomera martinezi | AMPHIBIA | 1,460 | 0.0052 | 12 | 4: Grassland |
| 78527890 | Oreophrynella seegobini | AMPHIBIA | 8 | 0.0012 | 12 | 3: Shrubland |
| 79101760 | Pseudophilautus puranappu | AMPHIBIA | 59 | 0.0002 | 300 | 3: Shrubland, 6: Rocky areas |
| 83410198 | Ctenophorus nguyarna | REPTILIA | 3,382 | 0.0002 | 9,007 | 5: Inland Wetlands |
| 83777277 | Varanus auffenbergi | REPTILIA | 1,275 | 0.0056 | 400 | 14.3: Plantations, 14.5: Urban |
| 83778246 | Varanus mertensi | REPTILIA | 1,078,556 | 0.0073 | 9,007 | 5: Inland Wetlands |
| 83778268 | Varanus mitchelli | REPTILIA | 610,034 | 0.0081 | 9,007 | 5: Inland Wetlands |
| 102325554 | Lerista vanderduysi | REPTILIA | 4,146 | 0.0018 | 9,007 | 3: Shrubland |
| 102345892 | Eutropis quadratilobus | REPTILIA | 15 | 0.0067 | 9,007 | 1: Forest |
| 103778254 | Orthotomus chaktomuk | AVES | 8,593 | 0.0021 | 100 | 3: Shrubland |
| 103874452 | Montecincla fairbanki | AVES | 4,283 | 0.0034 | 1,335 | 14.4: Rural Gardens, 3: Shrubland |
| 104681745 | Cnemaspis alantika | REPTILIA | 2,262 | 0.0044 | 250 | 1: Forest, 6: Rocky areas |
| 109452611 | Cryptoblepharus aldabrae | REPTILIA | 56 | 0.0030 | 12 | 14.4: Rural Gardens, Other |
| 109477411 | Lerista rochfordensis | REPTILIA | 92 | 0.0011 | 9,007 | 3: Shrubland |
| 109480639 | Carinascincus microlepidotus | REPTILIA | 20,015 | 0.0048 | 7,580 | 3: Shrubland, 6: Rocky areas |
| 110160538 | Hemicordylus nebulosus | REPTILIA | 22 | 0.0055 | 300 | 3: Shrubland, 6: Rocky areas |
| 112139816 | Rattus detentus | MAMMALIA | 1,900 | 0.0004 | 166 | 14.3: Plantations, 14.4: Rural Gardens, 14.6: Heavily Degraded Former Forest |
| 112685562 | Microkayla melanocheira | AMPHIBIA | 17 | 0.0053 | 159 | 3: Shrubland, 5: Inland Wetlands |
| 122894031 | Caledoniscincus pelletieri | REPTILIA | 67 | 0.0046 | 80 | 3: Shrubland |
| 127902191 | Sitana devakai | REPTILIA | 13,414 | 0.0083 | 9,007 | 3: Shrubland, Other |
| 137792366 | Calyptommatus leiolepis | REPTILIA | 7,456 | 0.0006 | 9,007 | 1: Forest |
| 181489412 | Anaxyrus nevadensis | AMPHIBIA | 19 | 0.0042 | 12 | 5: Inland Wetlands |

#### Validation using species' point observations

A total of 746 species had a true AOH map and sufficient data to use in point validation (426 birds, 137 mammals, 105 amphibians and 78 reptiles). Mean point prevalence was 0.95 (SD 0.15) overall and 0.96 (SD 0.15) for birds, 0.96 (SD 0.16) for mammals, 0.95 (SD 0.17) for amphibians and 0.93 (SD 0.18) for reptiles. A total of 730 species' AOH maps (98% of those included in point validation) performed better than expected if AOH were distributed within the range at random. This left 16 that had a lower point prevalence than map prevalence (6 birds, 5 mammals, 4 amphibians and 1 reptile)(Table S2.6).

**Table S2.6.** Details of the 16 species' AOH maps for which point prevalence was less than map prevalence indicating lower performance than expected if AOH were distributed within the range at random.

| TaxonID | Scientific name | Class | Range size (Number of cells) | Map prevalence | Elevation range (m) | Habitats |
|---|---|---|---|---|---|---|
| 977 | Alytes muletensis | AMPHIBIA | 182 | 0.01 | 840 | 14.2: Pasture, 15: Artificial Aquatic, 5: Inland Wetlands |
| 16750 | Petrogale xanthopus | MAMMALIA | 26,597 | 0.00 | 9,007 | 6: Rocky areas |
| 18267 | Procyon pygmaeus | MAMMALIA | 514 | 0.70 | 20 | 14.1: Arable, 1: Forest |
| 18557 | Pseudomys shortridgei | MAMMALIA | 32,071 | 0.02 | 9,007 | 3: Shrubland |
| 41656 | Mustela itatsi | MAMMALIA | 288,186 | 0.62 | 336 | 14.1: Arable, 14.2: Pasture, 14.3: Plantations, 14.4: Rural Gardens, 1: Forest, 3: Shrubland, 4: Grassland |
| 58662 | Lithobates megapoda | AMPHIBIA | 63,510 | 0.25 | 697 | 1: Forest, 3: Shrubland, 5: Inland Wetlands |
| 136113 | Rana draytonii | AMPHIBIA | 10,673 | 0.19 | 2,400 | 15: Artificial Aquatic, 1: Forest, 5: Inland Wetlands |
| 136509 | Petrogale mareeba | MAMMALIA | 20,729 | 0.00 | 1,000 | 6: Rocky areas |
| 157271 | Iberolacerta galani | REPTILIA | 1,004 | 0.00 | 900 | 6: Rocky areas |
| 22685505 | Psittacula alexandri | AVES | 2,431,048 | 0.88 | 2,000 | 14.1: Arable, 14.3: Plantations, 14.4: Rural Gardens, 14.5: Urban, 14.6: Heavily Degraded Former Forest, 1: Forest |
| 22693359 | Calidris tenuirostris | AVES | 50,357,714 | 0.17 | 1,300 | 4: Grassland, Other, Other, Other |

| TaxonID | Scientific name | Class | Range size (Number of cells) | Map prevalence | Elevation range (m) | Habitats |
|---|---|---|---|---|---|---|
| 22694053 | Vanellus gregarius | AVES | 7,803,638 | 0.36 | 300 | 14.1: Arable, 4: Grassland, 5: Inland Wetlands, 8: Desert |
| 22698230 | Puffinus yelkouan | AVES | 35,043 | 0.04 | 9,007 | 6: Rocky areas, Other, Other, Other |
| 22721040 | Calcarius ornatus | AVES | 3,318,748 | 0.44 | 1,500 | 3: Shrubland, 4: Grassland |
| 22721737 | Setophaga striata | AVES | 16,323,078 | 0.12 | 2,400 | 1: Forest, 3: Shrubland |
| 77318429 | Andinobates opisthomelas | AMPHIBIA | 7,855 | 0.47 | 750 | 1: Forest |

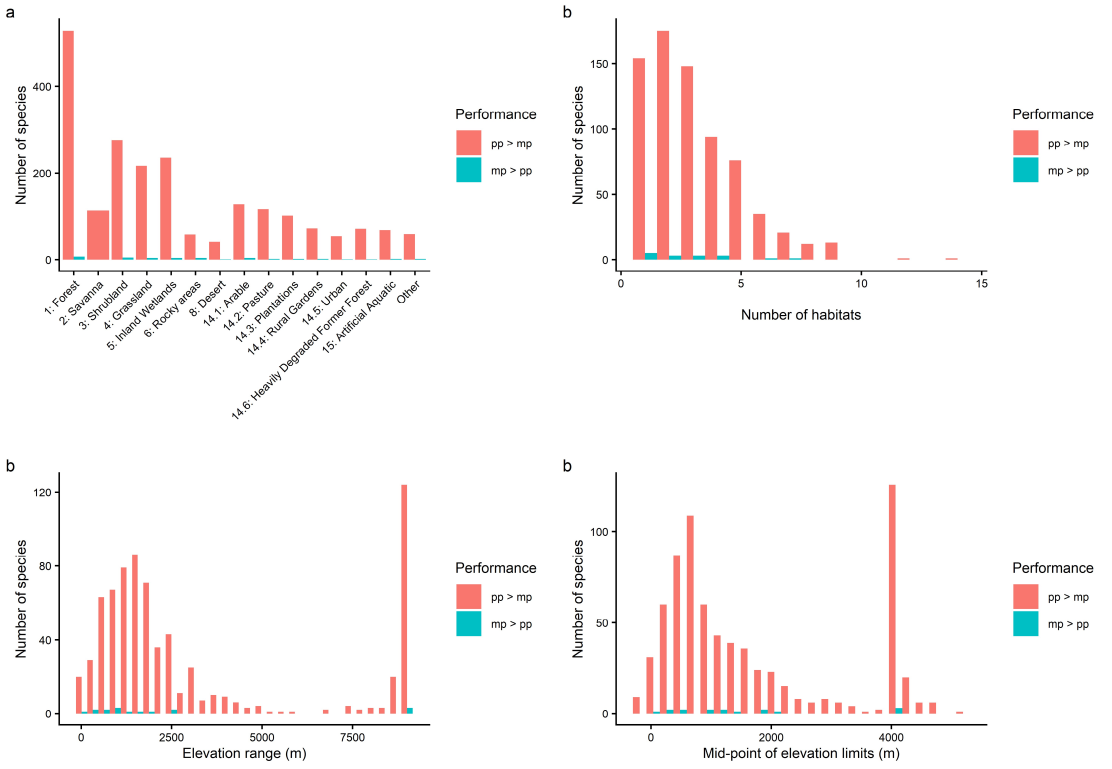


**Figure S2.3.** Comparison of species AOH maps with better than random performance with those with worse than random performance a) Habitats, b) number of habitats, c) elevation range, d) elevation mid-point.
