## Supplementary File 2 - Results and validation for "Global threat abatement potential for terrestrial vertebrates"

### Appendix 1. Supplementary methods and tables

**Adjustments and corrections to species' elevation associations data**

Errors and omissions in species' elevation associations were modified following (Baisero, 2021). Where either the upper or lower elevation limit was missing, or outside the range of global values present in the FABDEM data, the missing or incorrect limits were set to the global maximum (8580 m) or minimum (-427 m) FABDEM value respectively. Where the lower elevation limit was higher than the upper elevation limit, both upper and lower limits were replaced with the upper and lower limits of the FABDEM data respectively. Species' elevation limits were also adjusted for species that had an elevation range less than the vertical accuracy of FABDEM. If the elevation range was less than 12 m, both the upper and lower limits were widened to 6 m above or below the elevation mid-point respectively. These values were chosen based on the highest mean absolute error of FABDEM across different land-cover classes (Hawker et al., 2022).

For birds, elevation limits were supplemented with minimum and maximum 'occasional' elevation limits from BirdLife International (2024). These reflect extremes that are used in some years with atypical seasonal conditions.

**Adjustments to AOH**

If no suitable land-cover at suitable elevation was identified within the terrestrial range, adjustments to the AOH method were applied. If there was no suitable elevation within the range but there was some suitable land-cover then AOH was defined using suitable land-cover within the terrestrial range only. Conversely, if there was no suitable land-cover within the terrestrial range but there were cells at suitable elevation then AOH was defined using elevation within the terrestrial range only. If neither suitable land-cover nor suitable elevation was identified within the terrestrial range, the whole terrestrial range was considered to contain potentially suitable habitat. This step was taken as a precaution against omission errors.

**Correction to AOH for missing terrestrial land-cover over islands**

Validation revealed errors in the CGLS100 dataset whereby no terrestrial land-cover classes occurred on a considerable number of island states, overseas territories and offshore islands or islets, including French Polynesia, Federated states of Micronesia and Seychelles, with only partial coverage of terrestrial land-cover classes on others. For the 162 species whose entire range was affected by these errors, a correction was necessary to prevent them being falsely excluded from the calculation of $STAR_T$. It was not deemed suitable to assume the whole species' range, regardless of intersection with land, contained potentially suitable habitat as the species were not exclusively terrestrial and large commission errors over marine areas could be introduced. We first identified all Global Administrative Areas (GADM, version 4.1, https://gadm.org/data.html) that intersected the range of a species where no terrestrial land-cover classes were present and considered these administrative areas to be affected by the error. For species whose whole range was affected, the entire area of intersection between the species' range and affected administrative areas was assumed to be potentially suitable terrestrial habitat. A further 158 species were found to be affected by the error, but some AOH was successfully mapped elsewhere. For these species, the area of intersection between the species' range and any affected administrative zones was added to the existing AOH.

We acknowledge that this approach did not identify all islands affected by the error as it relied on species being endemic to affected areas and the correct alignment of spatial datasets. The range polygons of 28 species did not intersect any affected administrative areas or any CGLS100 terrestrial land-cover classes. These species were reported to the Red List authority and excluded. A total of 9,100 species remained, on which $STAR_T$ was calculated.

**Table S1.1** Expected percentage population decline over 10 years or three generations from the combinations of scope and severity scores per threat from Mair et al. (2021).

| | | Severity | | | | | |
|---|---|---|---|---|---|---|---|
| | | Very rapid declines (>30%) | Rapid declines (20-30%) | Slow, significant declines (<20%) | Negligible declines | No decline | Causing/could cause fluctuations |
| Scope | Whole (>90%) | 63 | 24 | 10 | 1 | 0 | 10 |
| | Majority (50-90%) | 52 | 18 | 9 | 0 | 0 | 9 |
| | Minority (<50%) | 24 | 7 | 5 | 0 | 0 | 5 |

**Table S1.2**. The number of species with missing or unknown scope and severity for at least one threat of all STAR-relevant species with at least one relevant threat.

| | Number of species relevant to STAR* | Missing or unknown scope and severity for at least one threat | |
|---|---|---|---|
| | | Number of species | % of total |
| Amphibian | 3214 | 3093 | 96 |
| Bird | 2143 | 825 | 38 |
| Mammal | 1685 | 1329 | 79 |
| Reptile | 2308 | 1910 | 83 |
| Total | 9350 | 7157 | 77 |

*Relevance was based on the species' IUCN Red List Assessments. These values include species that were later removed from $STAR_T$ if Area of Habitat (AOH) was unable to be mapped.

**Table S1.3**. The number of species-threat combinations where at least one of scope and severity was missing or unknown for all STAR-relevant species with at least one relevant threat.

| | Number of species relevant to STAR* | Missing or unknown scope and severity for at least one threat | |
|---|---|---|---|
| | | Number of species | % of total |
| Amphibian | 16054 | 14906 | 93 |
| Bird | 10088 | 1750 | 17 |
| Mammal | 8238 | 5287 | 64 |
| Reptile | 8015 | 5883 | 73 |
| Total | 42395 | 27826 | 66 |

*Relevance was based on the species' IUCN Red List Assessments. These values include species that were later removed from $STAR_T$ if Area of Habitat (AOH) was unable to be mapped.

**Species point occurrence data**

Species' point occurrence data from the IUCN Red List (version 2023-1), were supplemented with occurrences from the Global Biodiversity Information Facility (GBIF, Table S1.4). GBIF was searched for observations of species presence with coordinates of maximum 100 m uncertainty, recorded since 2012, with no geospatial issues flagged in the meta-data and excluding fossil and living specimens. Geospatial issues were defined according to the default occurrence download filter within the 'rgbif' package for R (Chamberlain et al. 2022). Point occurrence data for mammals and birds were additionally excluded if they were observed in 2018 or before as these points were used to train the model associating habitats coded on the IUCN Red List with land-cover classes (Lumbierres et al. 2022). Coordinates with fewer than three decimal places were removed, accounting for coordinates with trailing zeros that would result in false precision. Coordinates were additionally removed if they fell in seas or oceans, were exactly zero, had equal values for latitude and longitude, or coincided with country centroids, capital cities, or institutional addresses (implemented using the package CoordinateCleaner in R; Zizka et al. 2019). As the objective was to validate the estimation of AOH within the range not the range polygon itself, points were restricted to those that intersected the relevant species' range polygons. Points were randomly sampled to leave one point per species per 1 $km^2$ cell. Species with fewer than 10 remaining points in total were excluded from the validation using point locality data.

**Table S1.4.** Details of GBIF downloads

| | ID | Citation |
|---|---|---|
| Amphibian | 0060330-240506114902167 | GBIF Occurrence Download https://doi.org/10.15468/dl.a4dvrv Accessed from R via rgbif (https://github.com/ropensci/rgbif) on 2024-06-05 |
| Bird | 0060317-240506114902167 | GBIF Occurrence Download https://doi.org/10.15468/dl.gbzatb Accessed from R via rgbif (https://github.com/ropensci/rgbif) on 2024-06-05 |
| Mammal | 0060316-240506114902167 | GBIF Occurrence Download https://doi.org/10.15468/dl.w6xyn5 Accessed from R via rgbif (https://github.com/ropensci/rgbif) on 2024-06-05 |
| Reptile (non-testudine) | 0060314-240506114902167 | GBIF Occurrence Download https://doi.org/10.15468/dl.mcv865 Accessed from R via rgbif (https://github.com/ropensci/rgbif) on 2024-06-05 |
| Testudines | 0012226-241107131044228 | GBIF Occurrence Download https://doi.org/10.15468/dl.rtmpka Accessed from R via rgbif (https://github.com/ropensci/rgbif) on 2024-11-13 |

**Table S1.5**. Extent coordinates of panels b-e in figure 5 in the main text in Mollweide Equal Area Projection System (unit m).

| Panel | Region | Xmin, xmax | Ymin, ymax |
|---|---|---|---|
| B | South East Asia | 9001763, 16237988 | -1632916.6, 2627586.55 |
| C | Hawaii | -15365285, -14867884 | 2306971.6, 2733421.87 |
| D | Caribbean | -8727410, -5797494 | 903910.3, 3358494.40 |
| E | Galapagos islands | -9232125, -8925691 | -185682.9, 87335.53 |
